# An Unsupervised Machine Learning Paradigm for Artifact Removal from Electrodermal Activity in an Uncontrolled Clinical Setting

**DOI:** 10.1101/2021.10.22.465489

**Authors:** Sandya Subramanian, Bryan Tseng, Riccardo Barbieri, Emery N. Brown

## Abstract

**Objective:** Electrodermal activity (EDA) reflects sympathetic nervous system activity through sweating-related changes in skin conductance. To enable EDA data to be used robustly in clinical settings, we need to develop artifact detection and removal frameworks that can handle the types of interference experienced in clinical settings.

**Methods:** We collected EDA data from 69 subjects while they were undergoing surgery in the operating room. We then built an artifact removal framework using unsupervised learning methods and informed features to remove the heavy artifact that resulted from the use of surgical electrocautery during the surgery and compared it to other existing methods for artifact removal from EDA data.

**Results:** Our framework was able to remove the vast majority of artifact from the EDA data across all subjects with high sensitivity (94%) and specificity (90%). In contrast, existing methods used for comparison struggled to be sufficiently sensitive and specific, and none effectively removed artifact even if it was identifiable. In addition, the use of unsupervised learning methods in our framework removes the need for manually labeled datasets for training.

**Conclusion:** Our framework allows for robust removal of heavy artifact from EDA data in clinical settings such as surgery. Since this framework only relies on a small set of informed features, it can be expanded to other modalities such as ECG and EEG.

**Significance:** Robust artifact removal from EDA data is the first step to enable clinical integration of EDA as part of standard monitoring in settings such as the operating room.

## INTRODUCTION

Artifact detection and removal is required for any physiological data collection, especially in uncontrolled and ‘messy’ situations like in the hospital or at home. As sensors become more ubiquitous and optimized for comfort and convenience over signal quality, ensuring data quality is increasingly the responsibility of analysis algorithms that can quickly detect and correct artifact. Specifically, robust artifact removal is required for any physiological modality to become clinical standard, since artifact removal must be integrated into hardware systems to ensure high quality data for clinicians. Most of this artifact is clearly identifiable by eye and attributable to obvious sources such as patient movement, accidental removal or repositioning of sensors, or interference from other equipment [1]. However, automating what can be seen by eye can prove to be challenging. Common methods for artifact removal in simpler situations, such as thresholding, may not be sufficient for complex clinical environments. In addition, artifact rejection strategies must be optimized for minimal collateral damage in terms of removal of true data, especially in cases where temporal dependencies exist. Temporal dependencies may also warrant special considerations in methods development, for example favoring removal of multiple smaller chunks of data rather than a single continuous chunk.

Electrodermal activity (EDA) is one such physiological measure that is inexpensive and convenient to collect, but is not yet clinical standard because there are not rigorous tools to process and analyze it. EDA tracks the changing electrical conductance of the skin due to the activity of sweat glands, which are part of the body’s sympathetic ‘fight or flight’ reflex [2]. It has immense potential as a physiological marker to track sympathetic activation in situations such as pain or stress. Developing frameworks and methodologies to process it, including artifact detection and removal specific to clinical situations, would bring it one step closer to being used in the clinic.

Supervised learning tools have been used successfully in a number of clinical applications, including radiology and pathology [3]. However, in the case of artifact detection, creating a labeled training set is a non-trivial task that is not part of the clinical workflow. It would require significant manual labor to label each small increment of time as artifact or true data. Previous studies using advanced supervised machine learning methods, including deep learning, have relied on such expert labeled datasets [4-8]. The timescale of artifact is often a fraction of a second, so to minimize the amount of excess data labeled as artifact, the increments of time must be very small, increasing the manual labor of labeling. Different types of artifact may also require specific labeled training sets. Instead, unsupervised learning methods do not require labeled training sets, since they assign data to groups based on detecting patterns in the data. Given that artifactual data is easily identifiable by eye, it is reasonable to hypothesize that artifact is innately different from true data. Therefore, unsupervised methods should be able to differentiate between them given the appropriate features. In addition, they can detect more complex patterns that cannot be explicitly codified [9].

In this paper, we develop a pipeline for removing artifact from EDA using three unsupervised learning methods: isolation forest, K-nearest neighbor distance, and 1-class support vector machine (SVM). Specifically, we use EDA collected during surgery in the operating room, where there is maximal artifact due to interference from surgical cautery equipment. This is one of the most intense clinical situations, so by showing that we can robustly remove artifact in this scenario, we can demonstrate that our method is adaptable for any clinical situation, which moves EDA one step closer to being clinical standard. To feed into the unsupervised methods, we defined 12 features in half-second windows based on our own experimentation and guidance from existing literature.

EDA data were collected continuously during lower abdominal surgery in 69 human subjects. The source of most artifact was surgical cautery, which causes large visible deflections in the data every time it is turned on and off, which can be over 150 times in an average surgery at short, irregular intervals. Each time the cautery is turned on, it typically only remains on for a few seconds. While the cautery-induced deflections are clearly visible, to complicate matters, there are periods of intact by shifted (down typically) EDA between the deflections. Finally, the magnitude, sharpness, and direction of artifactual deflections vary across subjects. Fig. 1 shows a few example datasets with which include large artifacts.

**Fig. 1.**
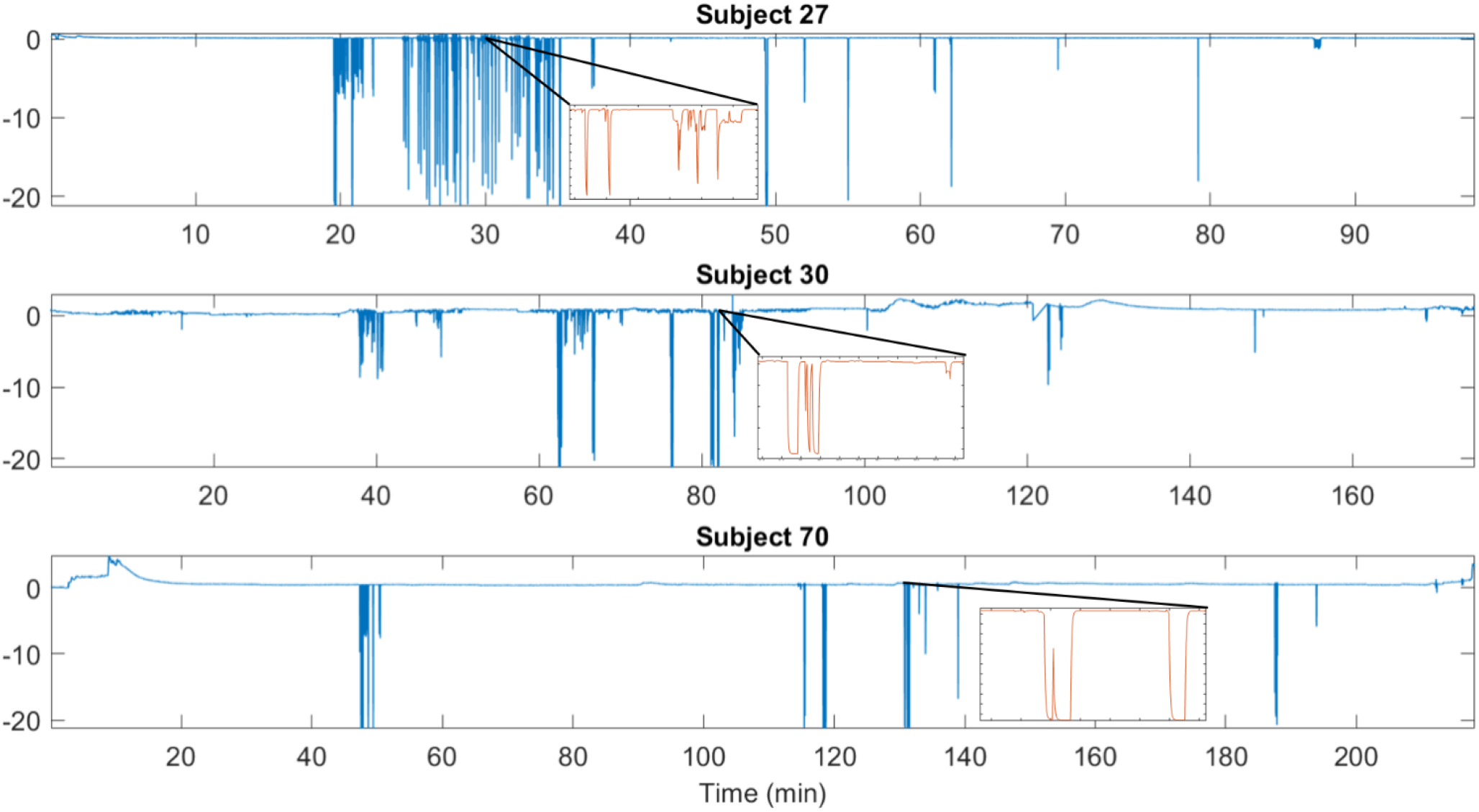
Examples of raw EDA data from three representative subjects

Existing unsupervised methods for artifact removal are specific to the datasets for which they were built, which were typically in mostly controlled experimental settings with occasional but minimal artifact [10-14]. None had the degree of artifact that surgical cautery interference produces. None were clinical EDA datasets. Unsurprisingly, neither variational mode decomposition nor wavelet decomposition, was able to successfully identify and remove the cautery-related artifact from clinical EDA data. In contrast, the artifact detection and removal pipeline we developed was able to successfully remove even heavy cautery artifact from all subjects’ data. In addition, our computational process was able to do so while preserving as much of the remaining true data as possible, including small snippets of real data in between sections of artifact. An earlier prototype of this methodology was published in [15].

In the remainder of this paper, we detail the development and validation of this pipeline. In Methods, we discuss the details of the data collection, subject cohort, the features used, and how the unsupervised learning algorithms were implemented to select artifact. In Results, we show each subject’s data before and after artifact removal and detail the specific parameters used and fraction of data labeled artifact. We also show a side-by-side comparison with existing artifact removal methods for a subset of the data. Finally, in Discussion, we address the implications of this work and our future directions.

## METHODS

### Data

In this study, we use EDA data recorded from 70 subjects (38 female, ages 29-77), collected under protocol approved by the Massachusetts General Hospital (MGH) Human Research Committee. All subjects were undergoing laparoscopic urologic or gynecologic surgery at MGH. The EDA data were recorded from two digits of each subject’s left hand at 256 Hz using the Thought Technology Neurofeedback System [16], starting from before induction of anesthesia to just after extubation. Fig. 1 shows an example of the raw data from three subjects. The main sources of artifact were movement at the beginning and end, including positioning, and use of surgical cautery. Each instance of turning cautery on or off caused a visible deflection in the data. Due to logistical concerns, EDA data collection from one subject (Subject 31) was ended before the onset of cautery, and therefore that subject was excluded from this analysis. EDA data from the remaining 69 subjects were analyzed using Matlab 2020b.

### Features and Unsupervised Learning Methods

The 12 features we used are listed in Table 1. These features are a combination of those used by other existing methods as well as additional ones that we discovered were useful based on experimentation. We computed these features for each 0.5 second window (128 samples) for each dataset to match with the timescale of most artifacts. These feature vectors were then fed as inputs into three unsupervised learning methods. Isolation forest is like random forest; however, each vector of features is scored based on the average length of the path to isolate it down to a leaf in an ensemble of decision trees [17]. Data that is artifact is thought to have a shorter path length than true data, since it is fundamentally different in nature from true data. In this case, each isolation forest consisted of 100 decisions trees, and the isolation scores were computed as the median of 10 such forests. K-nearest neighbor (KNN) distance computes the mean distance between each vector of features and the K nearest analogous vectors in the dataset [18]. In this case, KNN distance was computed using Euclidean distance and *K* = 50. Artifactual data is thought to be further in distance from true data. 1-class SVM is not unlike regular SVM, except that it is trained on only true data (only one class) and then tested on its ability to detect data that is not sufficiently similar to true data, in this case, artifact [19]. These artifacts are assumed to be rare in occurrence compared to true data. In this case, 1-class SVM was trained on 90% of the data, based on the 90% with the lowest KNN distance as a conservative estimate of true data and excluding the 10% of data points with the greatest KNN distance.

**Table 1.**
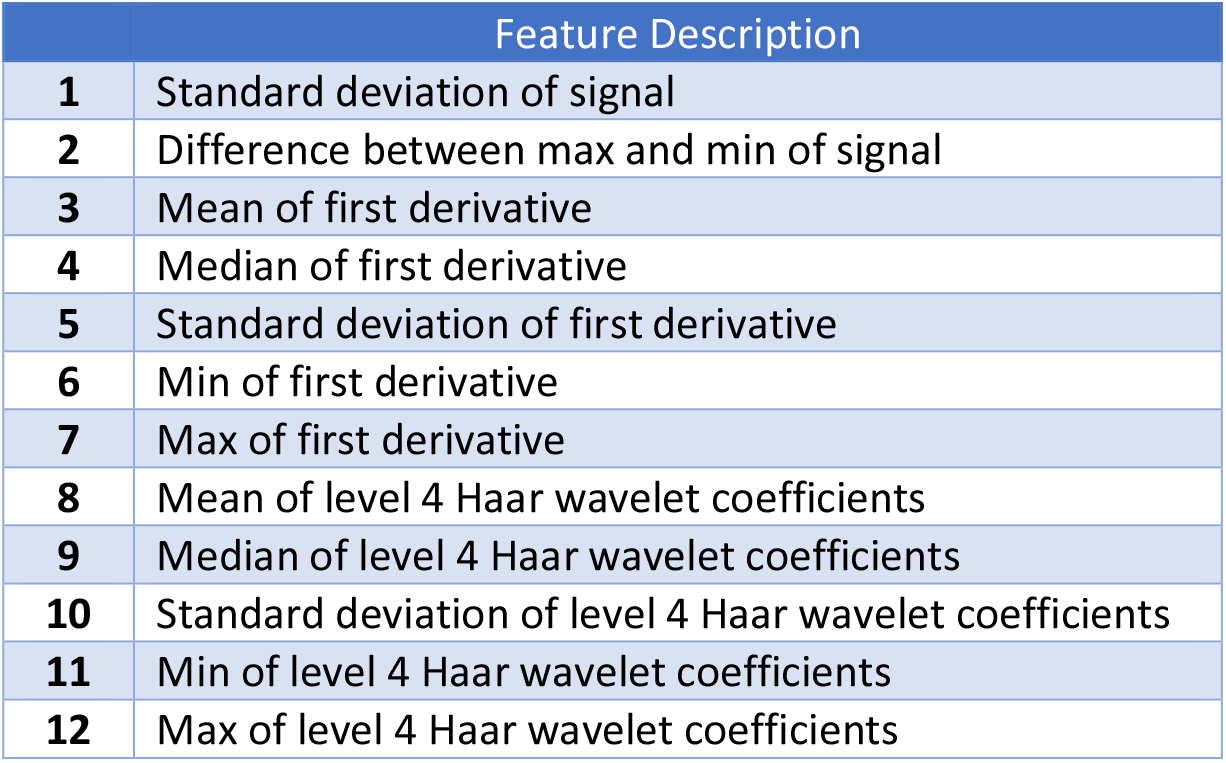
The 12 features for each 0.5 second window used as inputs for our unsupervised methods

All three unsupervised learning methods yielded a score for each window of data quantifying the degree of abnormality. The higher the score, the more likely that segment of data was artifact. The isolation forest scores were made negative to match the directionality of the other two. The last step of the process was to determine the appropriate threshold to determine artifact for each method for each subject. The process used to select these thresholds relied on specific insight about how all of the unsupervised methods label artifact. For each dataset, as the threshold on any of the unsupervised method scores is decreased, the portions of data that are labeled artifact increase in discrete jumps with more subtle changes in between. The most ‘correct’ labeling of artifact is likely to occur at one of these discrete jumps, since each jump represents the additional labeling of one similar ‘cluster’ of data as artifact whereas gradual changes represent a continuous spectrum of subtle differences within similar ‘clusters’. True artifact is highly similar to each other and distinctly different form true data; therefore, there should be no need to rely on subtle differences. To identify the discrete jump that represents the most ‘correct’ labeling of artifact, we took advantage of the fact that each discrete jump dramatically changes the inter-artifact interval distribution by introducing long gaps between subsequent artifact labels. Therefore, the skewness and kurtosis (3rd and 4th moments) of the inter-artifact interval distribution were computed across thresholds for each unsupervised method [20]. Since discrete jumps in labeled artifact skew the inter-artifact interval distribution, the jumps can be identified by local maxima in skewness and kurtosis. The thresholds at which local maxima in skewness and kurtosis occurred for each unsupervised method were tested by visually inspecting the labeled artifact. By using a binary search method to streamline which local maxima were tested, only 5-6 thresholds were visually inspected for each dataset. The best of these thresholds for each unsupervised method was selected by visual inspection. For each unsupervised learning method, once the ideal threshold was chosen, the total proportion of data labeled artifact was computed, as was the longest single continuous artifact.

After identifying and removing the artifact while preserving as much of the true data as possible, any ‘islands’ of true data that were shifted upward or downward due to artifactual deflection were translated back based on computing the linearly interpolated mean of the data at that time. The islands were typically clear in visual inspection, but quantitatively identifiable based on their minimum duration and minimum distance shifted up or down from the neighboring EDA data. The minimum duration and average distance were hyperparameters that were adjusted by subject to ensure no islands were excluded. After translating the ‘islands’ back, the gaps created by removed artifact were filled using linear interpolation once more to create continuous data. This is why the duration of the longest continuous artifact was relevant. Using linear interpolation to fill in a few seconds of data at a time will likely not affect downstream analysis; however, interpolating a few minutes at a time could.

Finally, we compared our method to other existing methods using both qualitative and quantitative methods. Using simple visual inspection, we compared our method to four other existing methods: variational mode decomposition [12-13], wavelet decomposition [10-11], thresholding at zero, and thresholding the derivative in terms of artifact removal. We also compared our method to two of those methods, wavelet decomposition and thresholding the derivative, in terms of precision of artifact detection. To do this, we randomly selected 21 subjects, and identified a 10-minute segment of data from each one. Twenty of the 10-minute segments were specifically selected to represent the regions of highest artifact for each respective subject, while the last segment was intentionally chosen to contain no artifact. Solely for the purposes of quantifying the performance of each method post-hoc, each 10-minute segment was divided into 0.5 second windows and manually labeled. These manual labels were used as ground truth to assess the classification performance of three methods that each have a distinct artifact detection step before an artifact removal step. For more details, see Supplementary Material Section S1. Variational mode decomposition does not have a separate artifact detection step and therefore could not be compared.

## RESULTS

Table S1 summarizes the results from all three unsupervised methods for all 69 subjects, including the final threshold chosen for each subject for each method. The proportion of data labeled artifact and the longest single continuous artifact are given (also summarized in Fig. 7). The best method, by smallest proportion of artifact removed (removing the least excess signal) and shortest maximum continuous artifact, is in bold for each subject. According to the proportion labeled artifact, isolation forest was the best method for 50 of the 69 subjects, KNN distance for 14 subjects, and 1-class SVM for 4 subjects, and both 1-class SVM and isolation forest were identical for one subject. Across all of the subjects, using isolation forest, the proportions of artifact ranged from 0.7% to just under 18% as shown in Fig. 7 and the longest contiguous artifact from 6 seconds to 194 seconds. For 52 of the 69 subjects (∼75%), the proportion labeled artifact was 10% or less and the longest continuous artifact was 30 seconds or shorter (Fig. 7).

Table S2 contains the hyperparameter values for identification of islands for each subject for all three unsupervised methods. Fig. 2 is a schematic summarizing our methodology for artifact removal using unsupervised learning. Fig. 3 shows an example of visually inspecting different thresholds for the same subject. All of the thresholds tested in Fig. 3 were chosen because they were local maxima of the skewness vs. threshold and kurtosis vs. threshold curves, as shown. The optimal threshold is determined by assessing degree of removal of artifact without unnecessary removal of signal. Fig. 4 shows the results after optimizing all three methods for artifact removal in three subjects. For one of the subjects (Subject 30), all three methods are able to similarly remove the artifact. For the other two subjects shown (Subjects 1 and 16), one or more of the methods are clearly superior to the others in removing the artifact without removing excess EDA signal. The uncorrected and final corrected EDA data using isolation forest, which was most often the best method, for all subjects are in shown in Figs S1 - S18 in the Supplementary Material. The degree of artifact varied across subjects, but we were able to remove the artifact in all cases. Figs. 5 and 6 show a comparison between our method and several existing methods for eight representative subjects in total. Both variational mode decomposition and wavelet decomposition were ineffective at artifact removal, and thresholding the EDA signal at 0 or thresholding the derivative of the EDA signal were partially effective though they still did not fully remove artifact. Only our method was effective to remove the majority of artifact. Table 2 exemplifies this further by showing the sensitivity and specificity of artifact detection using our method compared to wavelet decomposition and thresholding the derivative of the EDA signal on 10-minute segments of data from 21 randomly selected subjects. Overall, our method achieved 94% sensitivity and 90% specificity across all segments, whereas the other methods achieved either high sensitivity or high specificity, but not both. Our method was the only one that maintained above 50% sensitivity and specificity for all tested segments. In fact, for all but three segments of data, our method achieved 100% sensitivity and above 80% specificity. In contrast, wavelet decomposition results in many false positives, having overall low specificity (49%) while achieving high sensitivity (97%). In fact, in the segment of data with no artifact, wavelet decomposition still identifies over 80% artifact. In Figs. 5 and 6, while it appears that wavelet decomposition does not remove any artifact, this is due to ineffective artifact removal rather than artifact detection. On the other side, thresholding the derivative of the signal is not sensitive enough, achieving low overall sensitivity (46%) with high specificity (99%).

**Table 2.** Performance of different methods on 10-minute segments of data from 21 randomly selected subjects. Prop. artifact = proportion artifact, Sens - sensitivity, Spec = specificity

|  | Segment | Prop. artifact | Unsupervised learning + Informed features |  | Wavelet decomposition |  | Threshold derivative |  |
| --- | --- | --- | --- | --- | --- | --- | --- | --- |
|  |  |  | Sens | Spec | Sens | Spec | Sens | Spec |
| 1 | S16, 130-140 min | 0.0242 | 1 | 0.9752 | 1 | 0.4355 | 0.3793 | 1 |
| 2 | S65, 80-90 min | 0.2175 | 1 | 0.9712 | 1 | 0.1597 | 0.4559 | 0.9989 |
| 3 | S10, 155-165 min | 0.0358 | 1 | 0.9870 | 1 | 0.9844 | 0.3488 | 1 |
| 4 | S45, 105-115 min | 0.0742 | 1 | 0.8506 | 1 | 0.3240 | 0.5730 | 0.9946 |
| 5 | S7, 181-191 min | 0.1283 | 1 | 0.8824 | 1 | 0.7964 | 0.3442 | 1 |
| 6 | S58, 10-20 min | 0.0058 | 0.7143 | 0.9665 | 1 | 0.8927 | 0.4286 | 1 |
| 7 | S39, 70-80 min | 0.0725 | 1 | 0.8616 | 1 | 0.2956 | 0.4828 | 0.9856 |
| 8 | S68, 124-134 min | 0.0225 | 1 | 0.8142 | 1 | 0.5132 | 0.5185 | 0.9906 |
| 9 | S12, 105-115 min | 0.0383 | 1 | 0.9679 | 1 | 0.3752 | 0.5435 | 1 |
| 10 | S35, 50-60 min | 0.0567 | 1 | 0.9170 | 1 | 0.5150 | 0.4559 | 0.9956 |
| 11 | S57, 173-183 min | 0.0325 | 1 | 0.9854 | 1 | 0.3798 | 0.3333 | 0.9991 |
| 12 | S11, 165-175 min | 0.0583 | 1 | 0.9805 | 1 | 0.2451 | 0.4571 | 1 |
| 13 | S30, 60-70 min | 0.1300 | 0.5449 | 0.5766 | 0.9295 | 0.2510 | 0.2244 | 0.8688 |
| 14 | S66, 24-34 min | 0.0800 | 1 | 0.8904 | 1 | 0.4158 | 0.7396 | 0.9774 |
| 15 | S47, 160-170 min | 0.0100 | 1 | 0.9167 | 1 | 0.7795 | 0.6667 | 0.9975 |
| 16 | S3, 10-20 min | 0.0367 | 1 | 0.9965 | 1 | 0.5320 | 0.2727 | 1 |
| 17 | S61, 40-50 min | 0.0383 | 0.6304 | 0.6863 | 0.5217 | 0.3873 | 0.0870 | 0.9757 |
| 18 | S67, 15-25 min | 0.0675 | 1 | 0.9455 | 1 | 0.7712 | 0.7407 | 0.9821 |
| 19 | S49, 30-40 min | 0.0825 | 1 | 0.8265 | 1 | 0.2116 | 0.4545 | 0.9973 |
| 20 | S15, 110-120 min | 0.0333 | 1 | 0.8741 | 1 | 0.7931 | 0.7000 | 0.9948 |
| 21 | S70, 165-175 min | 0 | -- | 1 | -- | 0.1900 | -- | 1 |
|  | **AVERAGE** |  | **0.9445** | **0.8987** | **0.9726** | **0.4880** | **0.4603** | **0.9885** |

**Fig. 2.**
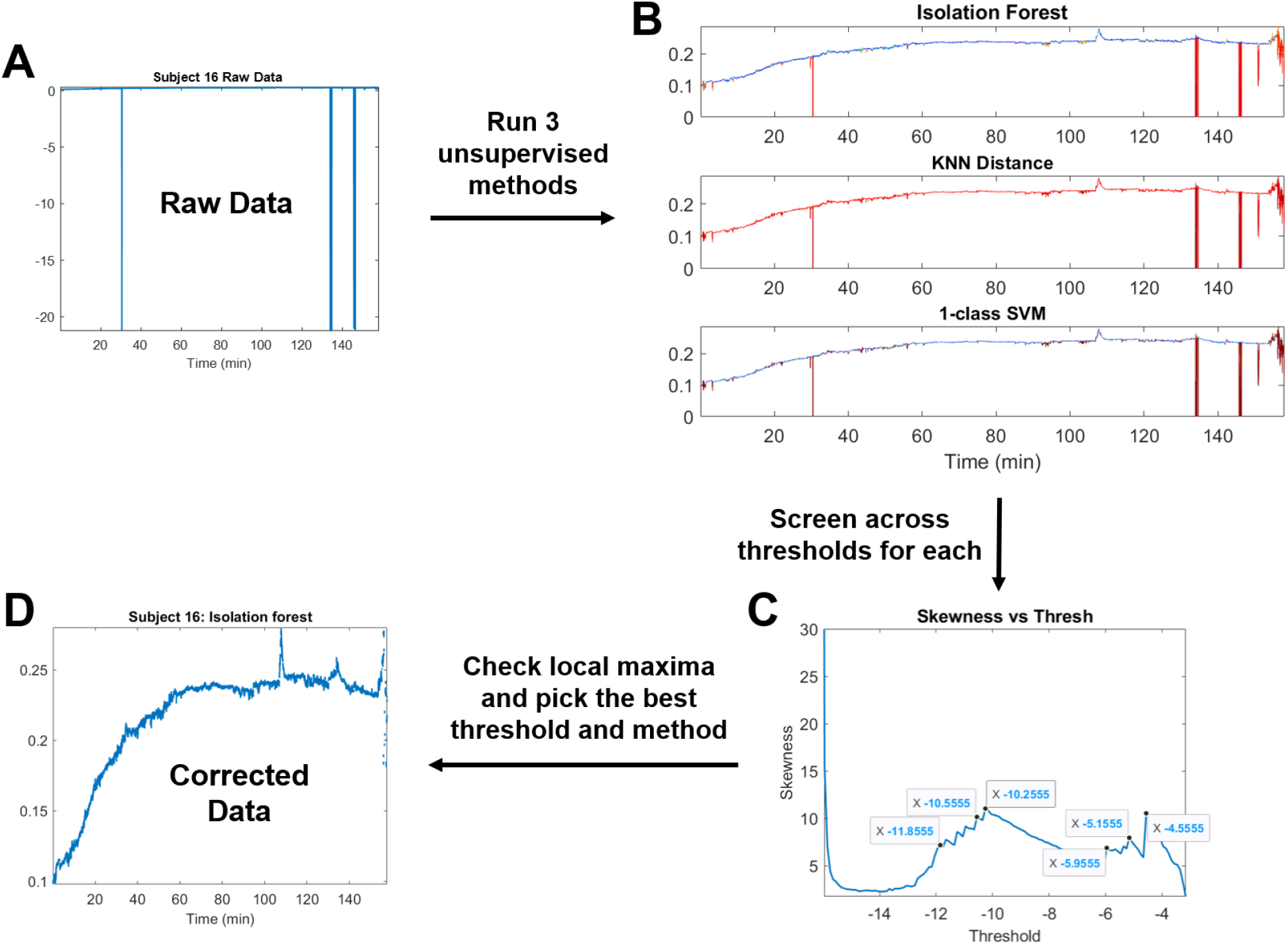
Schematic of methodology. (A) Starting from the raw EDA data, (B) each of the three unsupervised methods yields a score for each 0.5 second window of data, where a higher score is more likely artifact. (C) Screening across thresholds for artifact for all three scores, the labeled artifact at each threshold can be described by an associated inter-artifact interval distribution. The skewness and kurtosis of that distribution can be computed and plotted by threshold. (D) By visually inspecting the corrected EDA at the local maxima of the skewness vs. threshold and kurtosis vs. threshold curves, the best threshold for each of the three unsupervised methods can be selected. Then, the best method is the one which has the smallest proportion of labeled artifact.

**Fig. 3.**
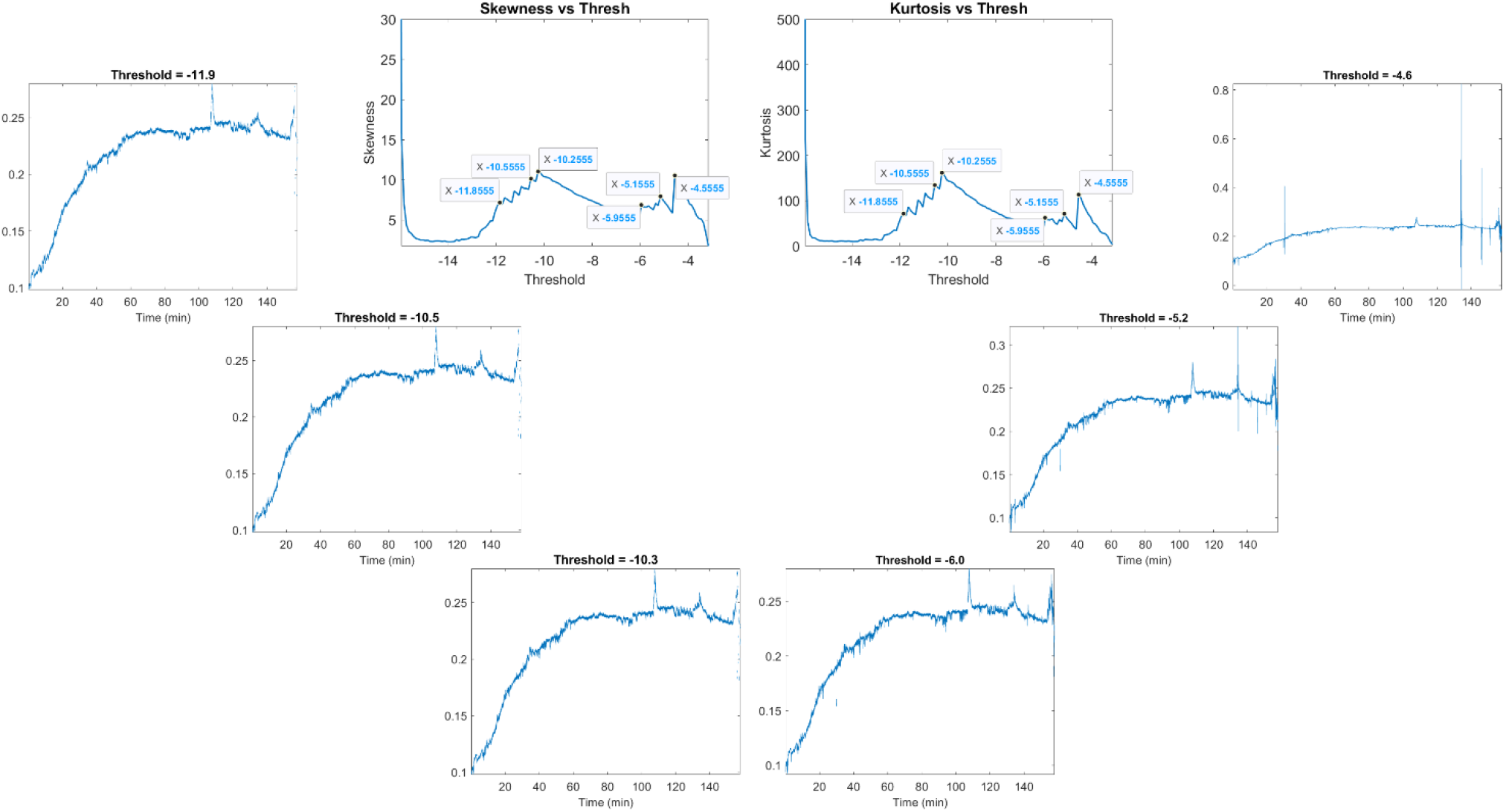
Comparing the artifact correction at multiple different local maxima of skewness and kurtosis for the same subject. The best threshold can be chosen by visually inspecting the corrected EDA at each of these thresholds and choosing the first threshold at which no artifacts are left behind without removing unnecessary excess signal.

**Fig. 4.**
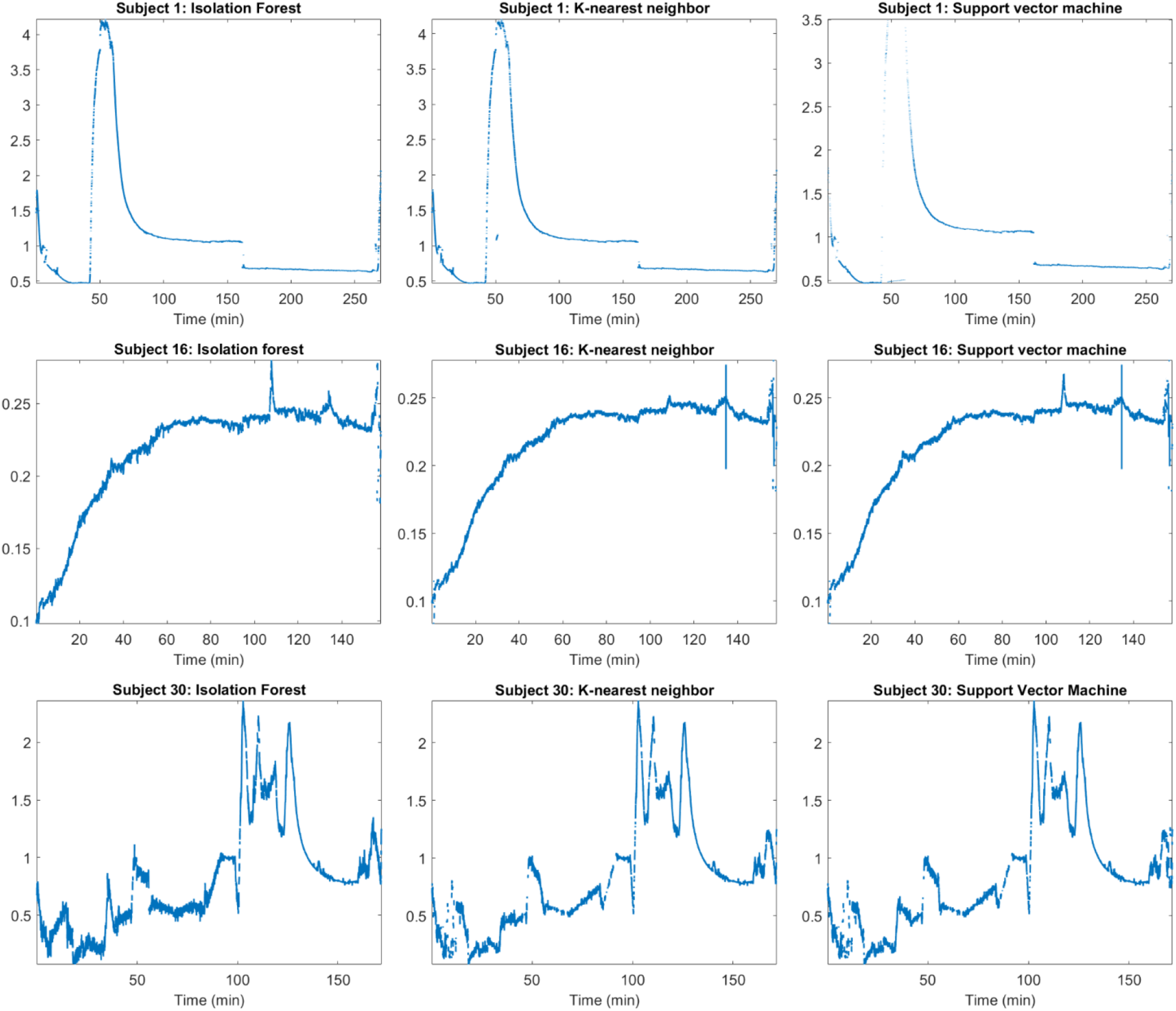
The results of artifact correction using each of the three unsupervised methods for three representative subjects. For some subjects, the different methods can all achieve similar performance (Subject 30), while for others, there are noticeable differences between the different methods (Subjects 1, 16).

**Fig. 5.**
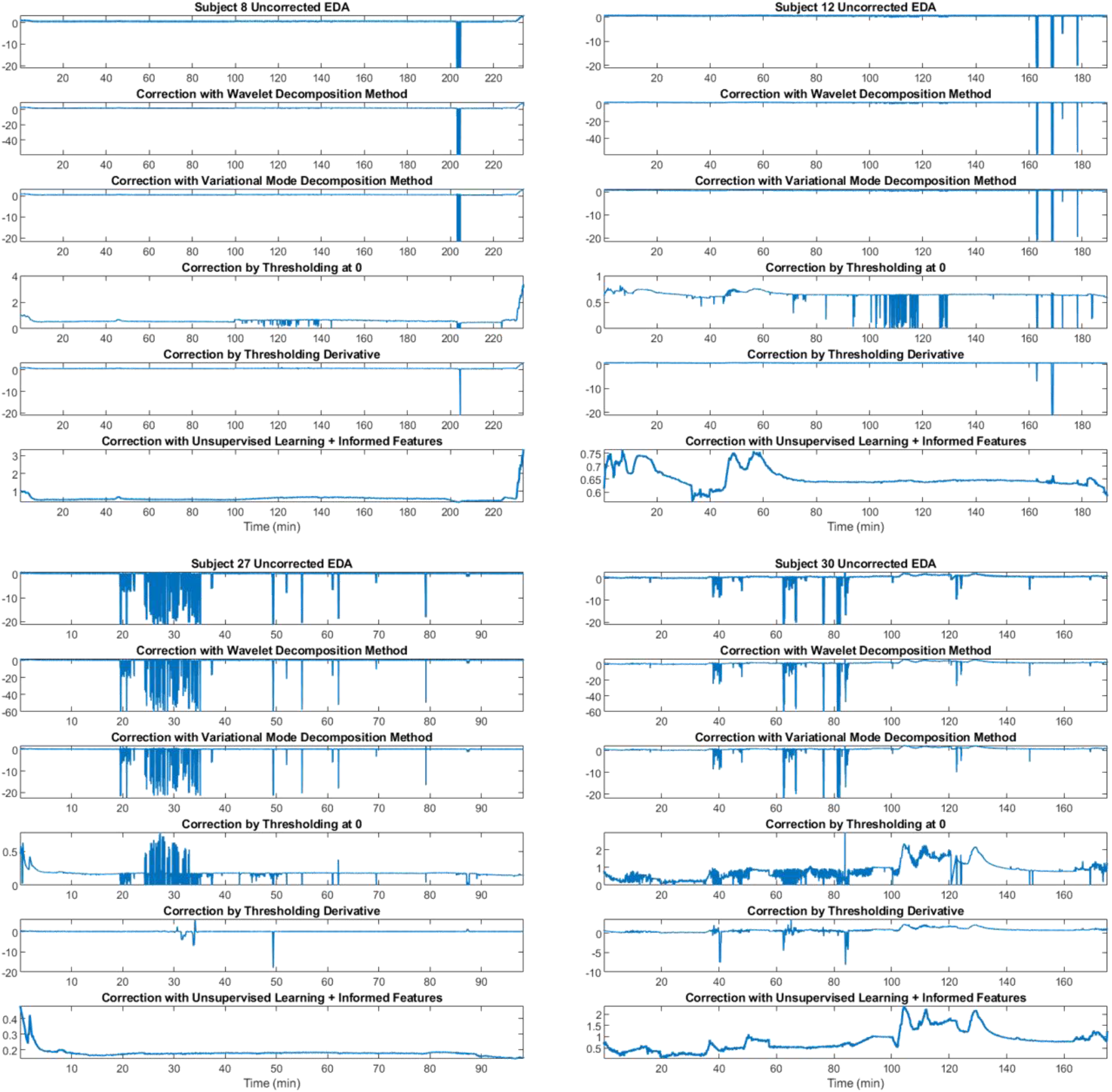
Comparison of artifact removal using unsupervised methods vs existing methods for 4 representative subjects

**Fig. 6.**
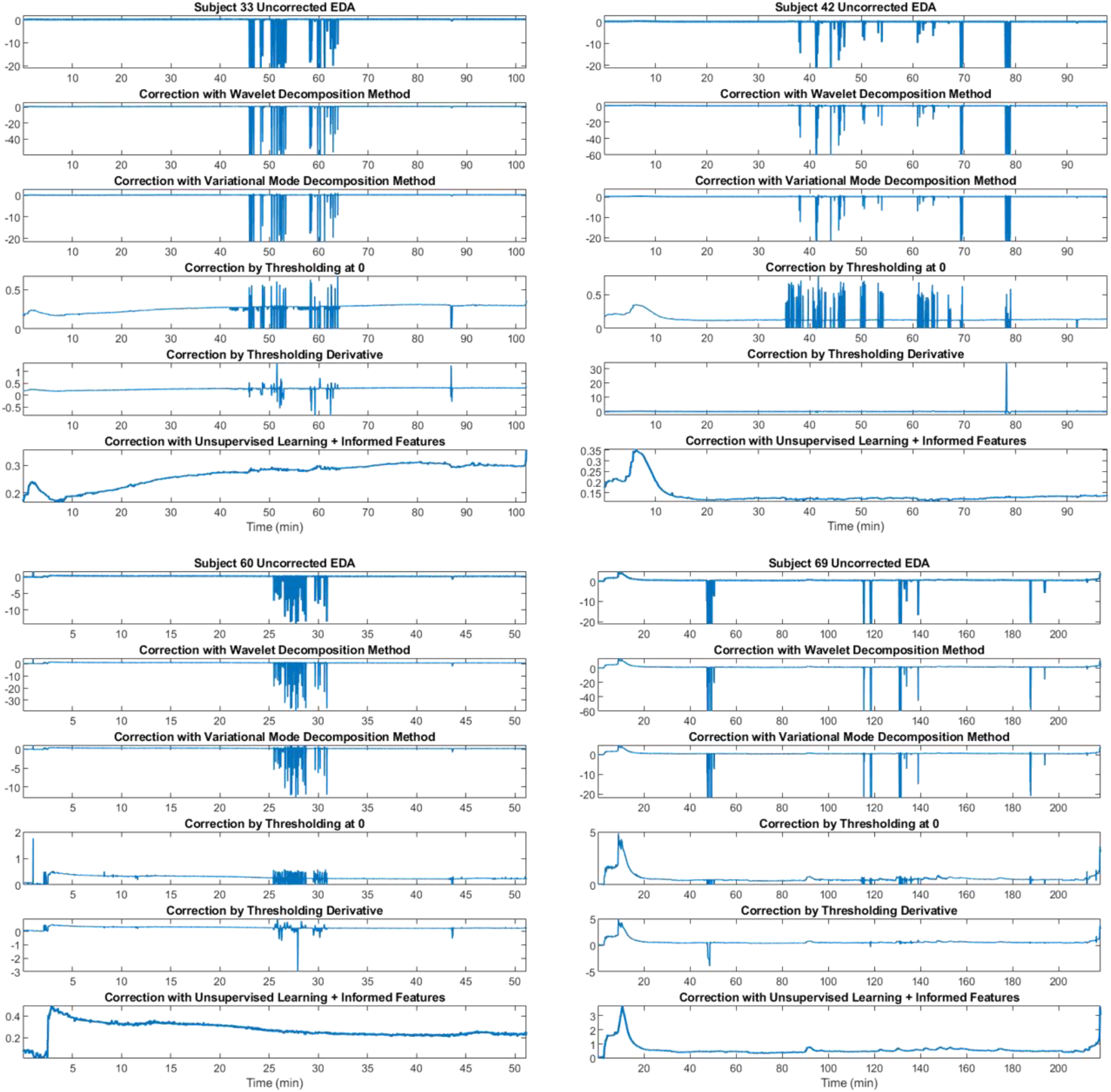
Comparison of artifact removal using unsupervised methods vs existing methods for 4 more representative subjects

**Fig. 7.**
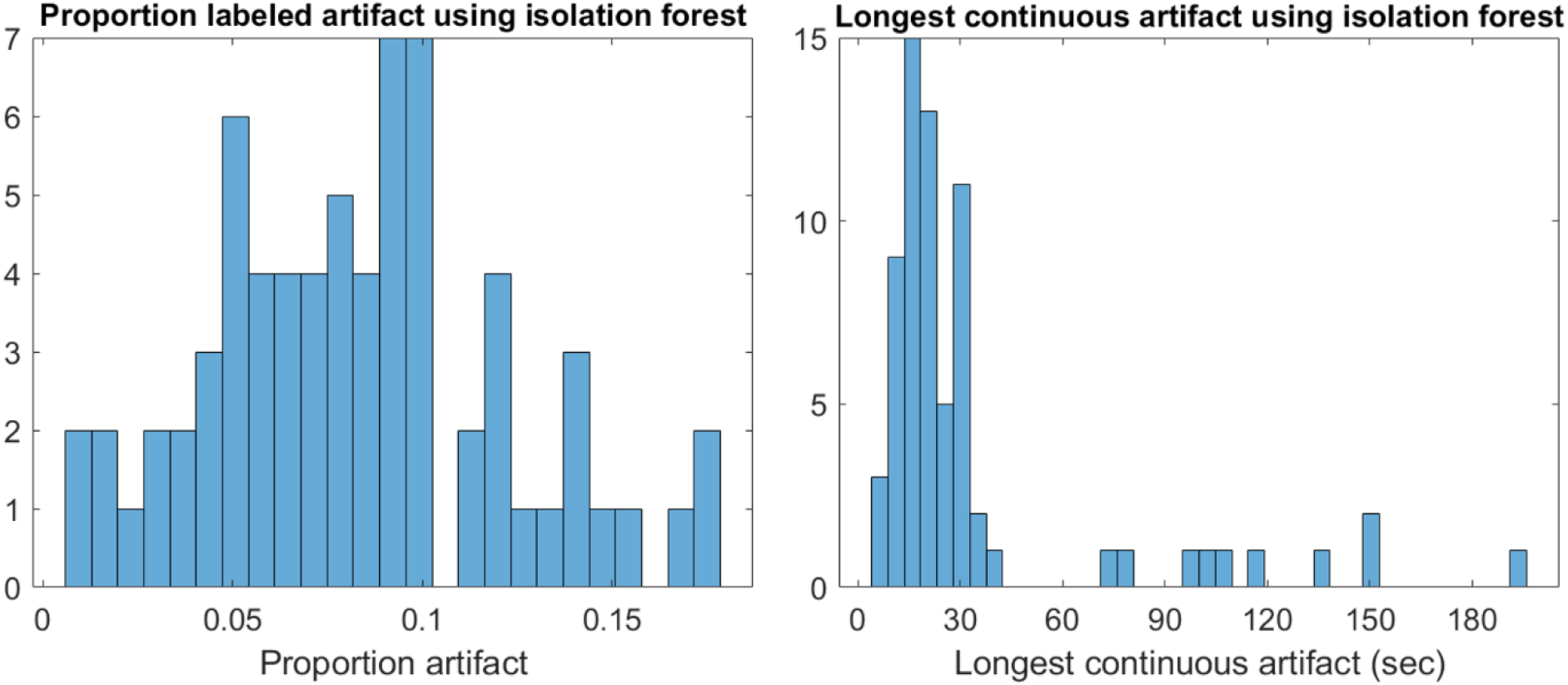
Distribution of total proportion of labeled artifact and longest continuous artifact across all 69 subjects

## DISCUSSION

In this study, we collected EDA data in the operating room during surgery from 69 subjects and demonstrated that using unsupervised machine learning methods and a set of 12 literature and physiology-informed features, we were able to remove artifact due to surgical cautery and movement from the EDA. This overcomes a major barrier for EDA to be used clinically. We specifically focused on unsupervised methods to emphasize practicality at the implementation stage, since a manually labeled training set is not required. We tested three unsupervised learning methods: isolation forest, K-nearest neighbor (KNN) distance, and 1-class support vector machine. We also compared existing methods such as variational mode decomposition and wavelet decomposition, as well as intuitive heuristic-based rules such as thresholding at zero or thresholding the derivative of the signal, for a representative subsample of subjects. Across all 69 subjects, the three unsupervised methods were able to remove the vast majority of artifact. None of the existing methods were able to fully remove major artifact in the tested subsample of subjects. Of the three unsupervised learning methods tested, isolation forest was the most discerning and parsimonious in not mislabeling true EDA as artifact for the majority of subjects (51 out of 69). Therefore, we used isolation forest for all subjects to arrive at the final artifact-removed EDA data.

Our methodology did not require any manual labeling of data for training, which would be extremely time-intensive and impractical in clinical settings. Despite the absence of training data, our methodology successfully removed cautery artifact from the data even when true EDA data was interspersed between sections of heavy artifact. This is indicated by the fact that even when the cautery seems to be continuous to the eye, the total proportion of labeled artifact in most cases (52 out of 69 subjects) was 10% or lower. In the subset of subjects in which comparison methods are shown in Figs 5 and 6, thresholding-based methods did not fully remove artifact; this was due to inadequate artifact detection for some methods and inefficient removal for others. Table 2 shows that the other methods were either not sensitive or specific enough; our method was the only one that maintained consistently high sensitivity and specificity across subjects. Even methods that could detect artifact were not necessarily able to remove it successfully (i.e. wavelet decomposition). Some of the comparison methods are decomposition-based, for example variational mode decomposition, which has the potential to affect the entire signal, including regions of true signal. In contrast, our method only modifies regions of the data that require modification and leaves non-artifact regions of data unchanged. In addition, most of the labeled artifact was in short segments of under 30 seconds. The longest continuous artifact only exceeded 60 seconds for 9 of the 69 subjects. This is important to consider in terms of downstream analysis, since relevant information about sympathetic activity is contained in the dynamic pulse-like phenomena in EDA [2,21]. Any method that modifies long, continuous chunks of data will likely affect the readout of dynamic activity in that timeframe. In contrast, short regions of missing data can be interpolated since they are only likely to contain a few pulses, and the missing data can be account for in estimation of uncertainty [22].

While our methodology used some of the same features as existing methods, we allowed the unsupervised algorithms to ‘learn’ the difference between artifact and true signal for each dataset on their own rather than hardcoding rules. The selected features, including those that overlap with existing methods, simply highlighted relevant characteristics of the data, based on the physiology of EDA and observations about the nature of cautery-related artifact. A straightforward expansion of this approach for other types of “clearly visible” artifact in modalities such as ECG and EEG could be implemented using custom feature definition, again informed by the physiology and nature of artifact in those signals. While our methodology is not fully automated at this stage and requires some visual examination for selection of hyperparameters, future iterations of our methodology will automate these steps as well.

## CONCLUSION

EDA data has great potential as a clinical marker of sympathetic activation; however, it is limited by the lack of hardware systems and software tools that have been built specifically for the clinical setting. This includes the crucial steps of artifact removal, specific to the degree and types of artifact that occur in clinical settings. Cautery interference during surgery is among the most intense of these, since there is an abundance of high-powered electrical equipment in use during surgery. Since most existing methods were built for purely experimental settings, none were effective in this scenario. However, we demonstrated that our paradigm is successfully able to recover viable EDA signal even in this situation, which allows EDA to be of clinical use as a marker of sympathetic activation even in the operating room. Future clinical EDA systems can integrate our methodology into their hardware to identify and remove artifacts as they occur. Our work is a critical advance to the eventual integration of EDA into clinical workflows as a biomarker of sympathetic activation, for example to track unconscious pain during surgery.

## Notes

### Competing Interest Statement

The authors have declared no competing interest.

