## Supplementary Material for "An Unsupervised Machine Learning Paradigm for Artifact Removal from Electrodermal Activity in an Uncontrolled Clinical Setting"

### Section S1. Detailed description of process for quantitative comparison between methods

**Selection of segments.** Twenty-one subjects were chosen at random without replacement from the full dataset. For the first twenty, the subject's EDA data were visually examined and a 10-minute segment was chosen from the period with highest density of artifact. For the last subject, a 10-minute segment without any artifact was chosen.

**Labeling ground truth.** Each segment was divided into non-overlapping half-second windows. Each window was visually examined and scored as 0, indicated true signal, or 1, indicated artifact. A window was scored as artifact if any portion of it contained artifact.

**Comparison methods**

Wavelet decomposition: Compute the level 4 Haar wavelet coefficients for the signal. Find the values that mark the $x$th percentile and $(100 - x)$th percentile of the detailed coefficients, zero out all detailed coefficients beyond those values in either direction. Any half-second window with zeroed coefficients was labeled as artifact. To aid the algorithm as much as possible, $x$ was set to half the true proportion of artifact. For artifact removal, the inverse wavelet transform would be performed with the new coefficients.

Thresholding the derivative: Compute the gradient and set the threshold for the absolute value of the derivative as 5. Anything above this is predicted to be artifact. Any half-second window contained predicted artifact was labeled as artifact.

### Section S2. Hyperparameter tables

**Table S1.** Thresholds, proportions labeled artifact, and longest single continuous artifacts for all 69 subjects. For each subject, the lowest proportion artifact and the shortest continuous artifact are in bold. SVM refers to 1-class support vector machine, IF refers to isolation forest, and KNN refers to K-nearest neighbor distance

| Subject | Threshold | | | Proportion Artifact | | | Longest continuous artifact (sec) | | |
|---|---|---|---|---|---|---|---|---|---|
| | SVM | IF | KNN | SVM | IF | KNN | SVM | IF | KNN |
| 1 | -323.07 | -7.88 | -0.4 | 0.1906 | **0.0418** | 0.077 | 55.0039 | **20.0039** | **20.0039** |
| 2 | -55.07 | -12.62 | -5.25 | 0.1144 | **0.0655** | 0.0843 | 41.5039 | **29.5039** | 41.0039 |
| 3 | -0.93 | -8.46 | -0.662 | 0.0599 | **0.0436** | 0.0564 | 45.5039 | **17.5039** | 45.5039 |
| 4 | -6.2 | -9.5 | -0.431 | 0.1316 | **0.1009** | 0.1292 | 38.5039 | **30.5039** | 37.0039 |
| 5 | -5.75 | -12.41 | -0.604 | **0.0583** | 0.0797 | 0.0659 | **21.5039** | **21.5039** | **21.5039** |
| 6 | -10 | -16.18 | -6.39 | 0.1774 | **0.1565** | 0.1709 | 352.0039 | **34.5039** | 352.0039 |
| 7 | -368.3 | -13.5 | -0.7 | 0.1581 | 0.1718 | **0.1155** | **18.5039** | **18.5039** | **18.5039** |
| 8 | -9.45 | -10.18 | -0.49 | **0.0601** | 0.0786 | 0.0729 | **17.5039** | **17.5039** | **17.5039** |
| 9 | -266.9 | -10.73 | -0.681 | **0.1154** | 0.1185 | 0.1328 | **16.5039** | **16.5039** | **16.5039** |
| 10 | -15.96 | -7.29 | -1.19 | 0.1011 | **0.0553** | 0.0936 | 118.5039 | **42.0039** | 118.5039 |
| 11 | -22.1 | -11.6 | -0.677 | **0.0629** | **0.0629** | 0.0643 | **21.5039** | **21.5039** | **21.5039** |
| 12 | -9.09 | -10.78 | -1.09 | 0.0583 | **0.0508** | 0.0572 | 32.0039 | **29.0039** | 32.0039 |

| | | | | | | | | | |
|---|---|---|---|---|---|---|---|---|---|
| 13 | -13.15 | -9.11 | -0.764 | 0.116 | **0.0507** | 0.1252 | 9.5039 | **9.0039** | 9.5039 |
| 14 | -37.446 | -14.96 | -6.49 | 0.1285 | **0.1014** | 0.1169 | 26.5039 | **18.5039** | 26.5039 |
| 15 | -11.328 | -11.82 | -1.4 | 0.096 | **0.0805** | 0.0949 | **19.5039** | **19.5039** | **19.5039** |
| 16 | -241.6 | -10.5 | -1.04 | 0.2073 | **0.0496** | 0.1705 | **20.0039** | **20.0039** | **20.0039** |
| 17 | -15.85 | -10.54 | -0.945 | 0.3232 | **0.1768** | 0.3352 | 23.5039 | **8.5039** | 35.5039 |
| 18 | -40.5 | -9.94 | -0.36 | 0.0928 | **0.0677** | 0.0775 | **14.0039** | **14.0039** | **14.0039** |
| 19 | -21.01 | -10.9 | -1.011 | 0.0747 | **0.069** | 0.0776 | **28.5039** | **28.5039** | **28.5039** |
| 20 | -10.47 | -8.349 | -0.689 | 0.1173 | 0.11099 | **0.1057** | 731.0039 | **138.0039** | 731.0039 |
| 21 | -109.4 | -12.77 | -1.126 | 0.0886 | **0.0697** | 0.0935 | 24.5039 | **21.5039** | 24.5039 |
| 22 | -400.9 | -8.85 | -0.565 | 0.1235 | **0.0319** | 0.1129 | **13.0039** | 79.1294 | **13.0039** |
| 23 | -352.5 | -8.9 | -0.452 | 0.096 | 0.05 | **0.0457** | **6.0039** | 72.6914 | **6.0039** |
| 24 | -5.36 | -8.58 | -0.097 | 0.0432 | **0.0365** | 0.0398 | **28.5039** | **28.5039** | **28.5039** |
| 25 | -7.1 | -11.12 | -0.849 | 0.0588 | **0.06** | 0.0614 | **28.5039** | **28.5039** | **28.5039** |
| 26 | -0.6 | -11.4 | -1.789 | 0.1997 | **0.0828** | 0.1269 | 28.0039 | **13.5039** | 18.0039 |
| 27 | -2.78 | -16.2 | -4.44 | 0.1434 | **0.1349** | 0.1362 | 46.0039 | **30.5039** | **30.5039** |
| 28 | -5.97 | -8.37 | -0.755 | 0.0814 | **0.0549** | 0.0789 | **23.5039** | **23.5039** | **23.5089** |
| 29 | -1.06 | -12.82 | -2.45 | 0.0906 | 0.1023 | **0.0812** | **40.5039** | 95.6914 | **40.5039** |
| 30 | -30.786 | -13.3 | -2.14 | 0.3188 | **0.2180** | 0.3154 | 103.5039 | **29.5039** | 103.5039 |
| 31 | No EDA | | | No EDA | | | No EDA | | |
| 32 | -63.8 | -12.73 | -1.08 | 0.1969 | **0.1215** | 0.1263 | **193.5039** | **193.5039** | **193.5039** |
| 33 | -11.67 | -12.7 | -3.41 | 0.0794 | **0.0563** | 0.0714 | 21.5039 | **16.0039** | 21.5039 |
| 34 | -4.86 | -12.4 | -1.34 | 0.1111 | **0.0977** | 0.1073 | 17.5039 | **16.0039** | 17.5039 |
| 35 | -10.05 | -13.65 | -2.82 | 0.1161 | **0.0935** | 0.1214 | 29.5039 | **24.5039** | 46.5039 |
| 36 | -9.5 | -16 | -3.39 | 0.1544 | **0.1484** | 0.154 | **22.5039** | **22.5039** | **22.5039** |
| 37 | -9.29 | -15.3 | -3.82 | 0.0986 | **0.0933** | 0.1058 | **13.5039** | **13.5039** | **13.5039** |
| 38 | -13.62 | -13.84 | -2.063 | 0.134 | **0.1172** | 0.1307 | **28.5039** | **28.5039** | **28.5039** |
| 39 | -10.69 | -15.42 | -4.32 | 0.0958 | **0.0879** | 0.0933 | **17.5039** | **17.5039** | **17.5039** |
| 40 | -6.53 | -13.37 | -2.518 | 0.0823 | **0.0762** | 0.0828 | 22.0039 | 22.0039 | **21.0039** |
| 41 | -258.57 | -10.1 | -0.497 | 0.1581 | **0.0425** | 0.0715 | **15.0039** | **15.0039** | **15.0039** |
| 42 | -2.6 | -14.14 | -3.3 | 0.0724 | **0.0696** | 0.0709 | 12.0039 | **11.5039** | **11.5039** |
| 43 | -55.3 | -13.01 | -1.13 | 0.124 | 0.1124 | **0.1032** | 12.5039 | 12.5039 | **12.0039** |
| 44 | -30.68 | -11.67 | -0.72 | 0.1083 | **0.0915** | 0.111 | 61.0039 | **31.5039** | 61.0039 |
| 45 | -8.3 | -16.2 | -3.26 | 0.1135 | 0.1245 | **0.1126** | **32.5039** | **32.5039** | **32.5039** |
| 46 | -9.5 | -13.14 | -1.83 | 0.1074 | **0.0934** | 0.1029 | **27.0039** | 100.0039 | **27.0039** |
| 47 | -13.5 | -6.4 | -0.32 | 0.0503 | **0.0234** | 0.0673 | **17.0039** | 20.0039 | **17.0039** |
| 48 | -89.5 | -8.69 | -0.67 | 0.1276 | **0.0512** | 0.0898 | 12.5039 | **11.0039** | **11.0039** |
| 49 | -2.1 | -13.25 | -2.466 | 0.1214 | **0.0946** | 0.1183 | 19.0039 | **18.0039** | **18.0039** |
| 50 | -3.69 | -13.1 | -1.794 | 0.1151 | **0.1002** | 0.1155 | **16.5039** | **16.5039** | **16.5039** |
| 51 | -118 | -11.86 | -1.08 | 0.1034 | **0.0911** | 0.1187 | **19.0039** | **19.0039** | 19.5039 |
| 52 | -3.96 | -13.34 | -4.0 | 0.0943 | 0.094 | **0.0888** | **21.0039** | 115.316 | **21.0039** |
| 53 | -24.7 | -12.34 | -0.973 | 0.1913 | **0.1382** | 0.1805 | **32.5039** | **32.5039** | **32.5039** |
| 54 | -4.02 | -11.99 | -1.635 | 0.1051 | 0.0869 | **0.0825** | 155.5039 | 34.5039 | **26.0039** |
| 55 | -2.4 | -9.93 | -0.2 | 0.0315 | 0.0309 | **0.029** | **15.5039** | **15.5039** | **15.5039** |
| 56 | -29.8 | -15.67 | -5.18 | 0.1557 | **0.1435** | 0.1448 | **24.5039** | 148.9409 | **24.5039** |
| 57 | -233.19 | -8.92 | -0.467 | 0.0764 | **0.0352** | 0.1389 | 24.0039 | **23.5039** | 24.5039 |
| 58 | -158.75 | -7.0 | 1.283 | 0.0392 | 0.0115 | **0.0044** | 12.0039 | 12.0039 | **11.5039** |
| 59 | -72.1 | -7.4 | 0.115 | 0.0278 | 0.0182 | **0.0129** | **10.0039** | 12.0039 | **10.0039** |
| 60 | -5.64 | -13.72 | -1.33 | 0.1028 | 0.1011 | **0.0974** | **14.5039** | **14.5039** | **14.5039** |

| | | | | | | | | | |
|---|---|---|---|---|---|---|---|---|---|
| 61 | -24.77 | -10.82 | -0.368 | 0.0766 | **0.0659** | 0.0837 | 19.5039 | **17.0659** | 19.5039 |
| 62 | -95.2 | -11.34 | -0.7 | 0.135 | **0.0527** | 0.0775 | 21.0039 | **13.5039** | **13.5039** |
| 63 | -88.46 | -13.07 | -1.03 | 0.1548 | **0.1179** | 0.1345 | 24.5039 | **23.6294** | 24.0039 |
| 64 | -66.6 | -6.45 | -0.06 | 0.0142 | **0.0066** | 0.0225 | 13.5039 | **6.0039** | **6.0039** |
| 65 | -19.745 | -13.77 | -4.07 | 0.0789 | **0.0741** | 0.0755 | **23.5039** | **23.5039** | **23.5039** |
| 66 | -12.35 | -13.34 | -0.885 | 0.1971 | 0.169 | **0.167** | 29.5039 | **14.5039** | 28.5039 |
| 67 | -5.46 | -12.2 | -1.47 | 0.0975 | **0.08** | 0.1323 | **8.0039** | **8.0039** | 10.5039 |
| 68 | -7.27 | -11.29 | -0.336 | 0.133 | 0.1399 | **0.1069** | 298.5039 | 151.0039 | **26.0039** |
| 69 | -56.15 | -12.29 | -0.877 | 0.0979 | **0.0853** | 0.0988 | 17.5039 | **14.0039** | 16.0039 |
| 70 | -60.06 | -15.22 | -4.74 | 0.1161 | **0.1002** | 0.1033 | 20.5039 | 20.5039 | **20.0039** |

**Table S2.** Hyperparameters for identification of 'islands' after each of the three unsupervised learning methods for all 69 subjects. SVM refers to 1-class support vector machine, IF refers to isolation forest, and KNN refers to K-nearest neighbor distance

| Subject | 1-class SVM | | Isolation Forest (IF) | | KNN Distance | |
|---|---|---|---|---|---|---|
| | Duration | Distance | Duration | Distance | Duration | Distance |
| 1 | 62 | 3 | 62 | 3 | 62 | 3 |
| 2 | 10 | 0.004 | 10 | 0.01 | 10 | 0.005 |
| 3 | 10 | 0.008 | 10 | 0.008 | 10 | 0.008 |
| 4 | 15 | 0.005 | 15 | 0.005 | 15 | 0.005 |
| 5 | 15 | 0.001 | 15 | 0.001 | 15 | 0.001 |
| 6 | 15 | 0.004 | 15 | 0.01 | 15 | 0.008 |
| 7 | 5 | 0.005 | 5 | 0.005 | 5 | 0.005 |
| 8 | 10 | 0.01 | 15 | 0.01 | 10 | 0.01 |
| 9 | 22 | 0.008 | 2 | 0.01 | 22 | 0.008 |
| 10 | 7 | 0.001 | 7 | 0.001 | 7 | 0.002 |
| 11 | 15 | 0.003 | 15 | 0.003 | 15 | 0.003 |
| 12 | 10 | 0.004 | 10 | 0.005 | 10 | 0.004 |
| 13 | 10 | 0.01 | 10 | 0.01 | 10 | 0.01 |
| 14 | 15 | 0.006 | 15 | 0.01 | 15 | 0.008 |
| 15 | 15 | 0.006 | 15 | 0.003 | 15 | 0.006 |
| 16 | 7 | 0.002 | 7 | 0.001 | 7 | 0.002 |
| 17 | 4 | 0.003 | 4 | 0.003 | 4 | 0.005 |
| 18 | 10 | 0.01 | 10 | 0.01 | 10 | 0.01 |
| 19 | 15 | 0.003 | 15 | 0.003 | 15 | 0.003 |
| 20 | 9 | 0.0025 | 9 | 0.0025 | 9 | 0.0025 |
| 21 | 15 | 0.002 | 15 | 0.003 | 15 | 0.002 |
| 22 | 10 | 0.015 | 10 | 0.022 | 10 | 0.02 |
| 23 | 15 | 0.01 | 15 | 0.01 | 15 | 0.01 |
| 24 | 15 | 0.003 | 15 | 0.003 | 15 | 0.003 |
| 25 | 15 | 0.003 | 15 | 0.003 | 15 | 0.003 |
| 26 | 5 | 0.005 | 5 | 0.005 | 10 | 0.005 |
| 27 | 19 | 0.005 | 19 | 0.005 | 8 | 0.005 |
| 28 | 20 | 0.01 | 20 | 0.01 | 20 | 0.01 |
| 29 | 20 | 0.01 | 20 | 0.01 | 20 | 0.01 |
| 30 | 15 | 0.01 | 15 | 0.02 | 15 | 0.01 |
| 31 | No EDA | | No EDA | | No EDA | |

| | | | | | | |
|---|---|---|---|---|---|---|
| 32 | 10 | 0.006 | 10 | 0.007 | 10 | 0.007 |
| 33 | 16 | 0.005 | 16 | 0.005 | 16 | 0.005 |
| 34 | 15 | 0.005 | 15 | 0.007 | 15 | 0.005 |
| 35 | 16 | 0.01 | 15 | 0.01 | 16 | 0.01 |
| 36 | 16 | 0.01 | 16 | 0.01 | 16 | 0.01 |
| 37 | 16 | 0.01 | 16 | 0.01 | 16 | 0.006 |
| 38 | 15 | 0.008 | 15 | 0.01 | 15 | 0.008 |
| 39 | 14 | 0.005 | 12 | 0.005 | 14 | 0.005 |
| 40 | 20 | 0.004 | 16 | 0.004 | 16 | 0.004 |
| 41 | 10 | 0.01 | 10 | 0.02 | 10 | 0.02 |
| 42 | 10 | 0.01 | 10 | 0.01 | 10 | 0.01 |
| 43 | 10 | 0.01 | 10 | 0.01 | 10 | 0.01 |
| 44 | 15 | 0.005 | 15 | 0.01 | 10 | 0.01 |
| 45 | 15 | 0.01 | 15 | 0.01 | 15 | 0.01 |
| 46 | 19 | 0.005 | 20 | 0.005 | 19 | 0.005 |
| 47 | 19 | 0.01 | 19 | 0.01 | 19 | 0.01 |
| 48 | 10 | 0.005 | 10 | 0.01 | 10 | 0.005 |
| 49 | 10 | 0.01 | 10 | 0.01 | 10 | 0.01 |
| 50 | 24 | 0.005 | 24 | 0.005 | 24 | 0.005 |
| 51 | 15 | 0.01 | 15 | 0.01 | 15 | 0.01 |
| 52 | 12 | 0.01 | 10 | 0.01 | 12 | 0.01 |
| 53 | 10 | 0.01 | 5 | 0.01 | 10 | 0.01 |
| 54 | 15 | 0.005 | 10 | 0.005 | 10 | 0.005 |
| 55 | 15 | 0.05 | 15 | 0.05 | 15 | 0.05 |
| 56 | 7 | 0.03 | 7 | 0.03 | 7 | 0.04 |
| 57 | 15 | 0.01 | 15 | 0.01 | 12 | 0.02 |
| 58 | 10 | 0.01 | 10 | 0.02 | 10 | 0.02 |
| 59 | 10 | 0.01 | 10 | 0.01 | 10 | 0.01 |
| 60 | 6 | 0.01 | 6 | 0.01 | 6 | 0.01 |
| 61 | 10 | 0.025 | 10 | 0.01 | 10 | 0.01 |
| 62 | 5 | 0.01 | 10 | 0.01 | 5 | 0.01 |
| 63 | 10 | 0.01 | 15 | 0.01 | 10 | 0.01 |
| 64 | 5 | 0.05 | 5 | 0.05 | 5 | 0.05 |
| 65 | 15 | 0.05 | 15 | 0.05 | 15 | 0.04 |
| 66 | 7 | 0.01 | 7 | 0.01 | 10 | 0.01 |
| 67 | 5 | 0.01 | 6 | 0.01 | 5 | 0.01 |
| 68 | 5 | 0.01 | 10 | 0.01 | 10 | 0.01 |
| 69 | 10 | 0.01 | 10 | 0.01 | 10 | 0.01 |
| 70 | 10 | 0.01 | 10 | 0.01 | 10 | 0.01 |

### Section S3. Additional figures

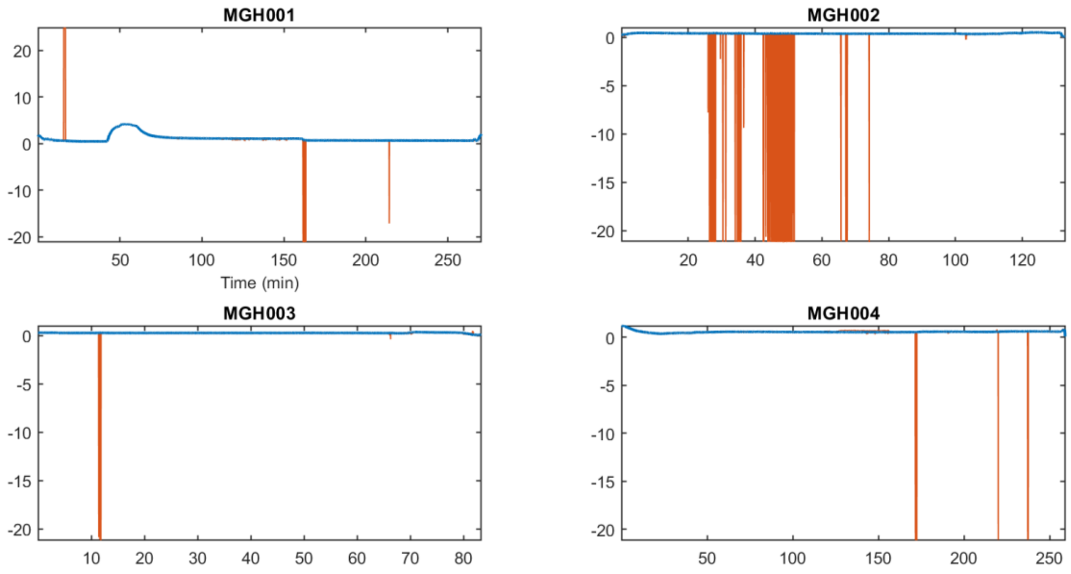


**Figure S1:** Raw and artifact-free EDA data for Subjects 1-4 collected while undergoing surgery at Massachusetts General Hospital (MGH). The raw data is in orange and the artifact-free data is in blue.

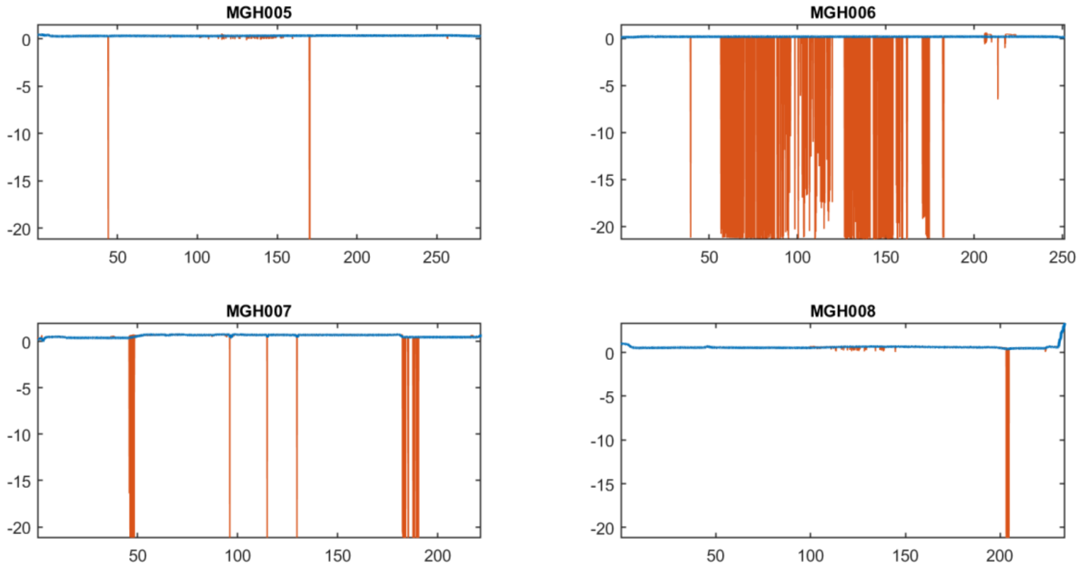


**Figure S2:** Raw and artifact-free EDA data for Subjects 5-8 collected while undergoing surgery at Massachusetts General Hospital (MGH). The raw data is in orange and the artifact-free data is in blue.

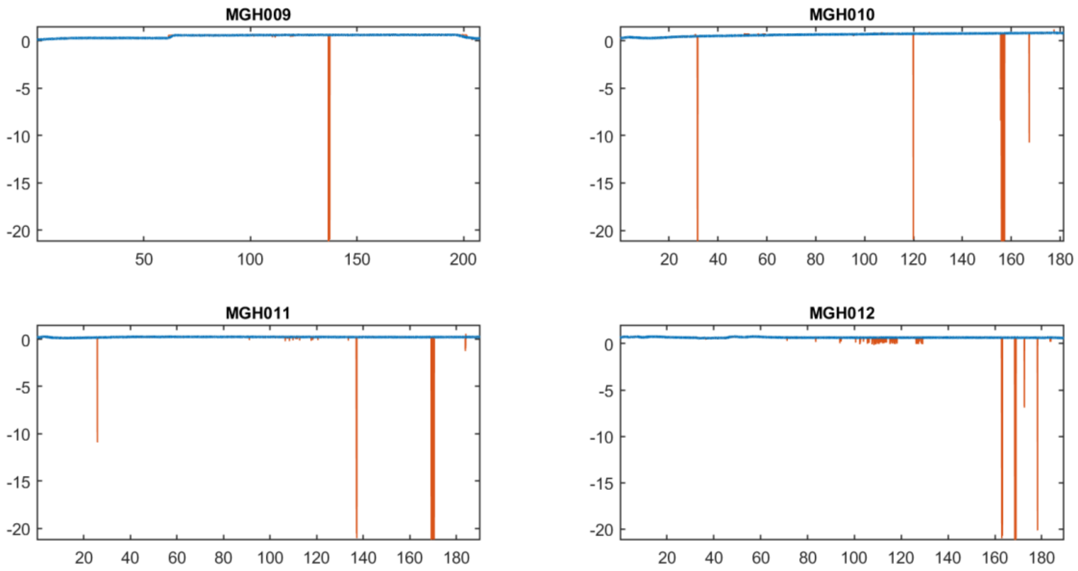


**Figure S3:** Raw and artifact-free EDA data for Subjects 9-12 collected while undergoing surgery at Massachusetts General Hospital (MGH). The raw data is in orange and the artifact-free data is in blue.

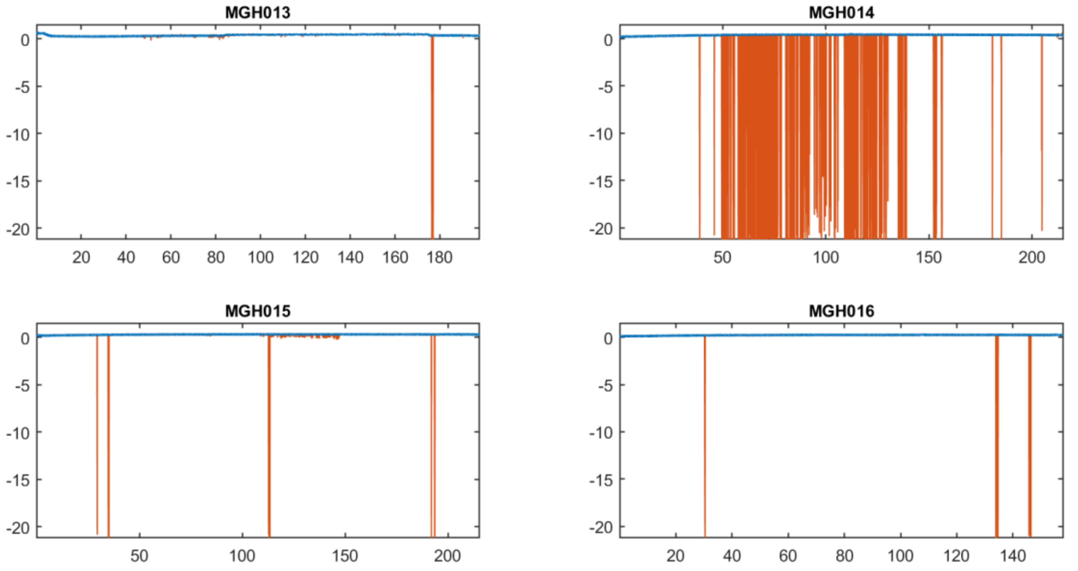


**Figure S4:** Raw and artifact-free EDA data for Subjects 13-16 collected while undergoing surgery at Massachusetts General Hospital (MGH). The raw data is in orange and the artifact-free data is in blue.

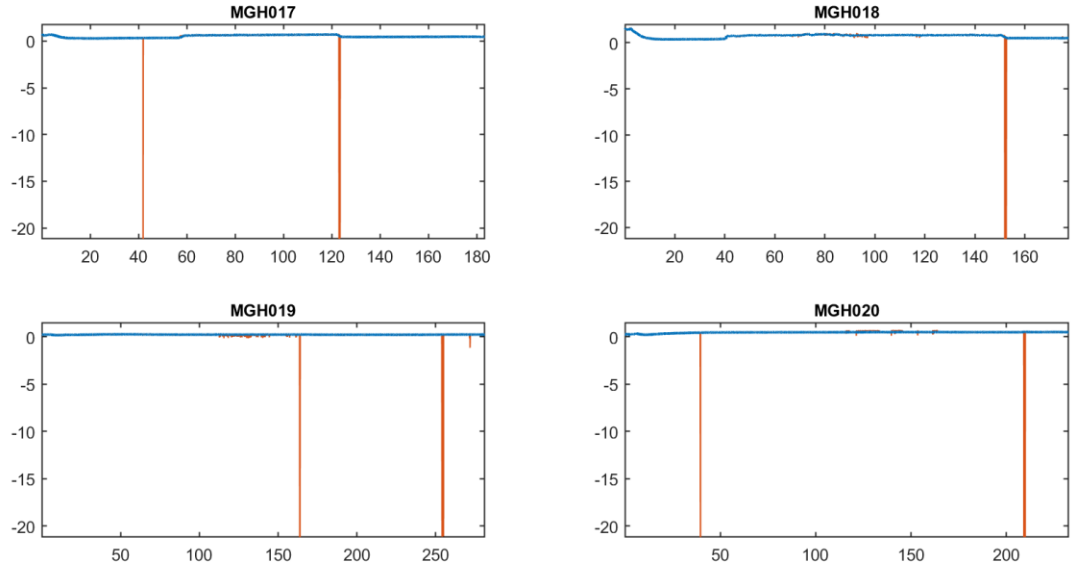


**Figure S5:** Raw and artifact-free EDA data for Subjects 17-20 collected while undergoing surgery at Massachusetts General Hospital (MGH). The raw data is in orange and the artifact-free data is in blue.

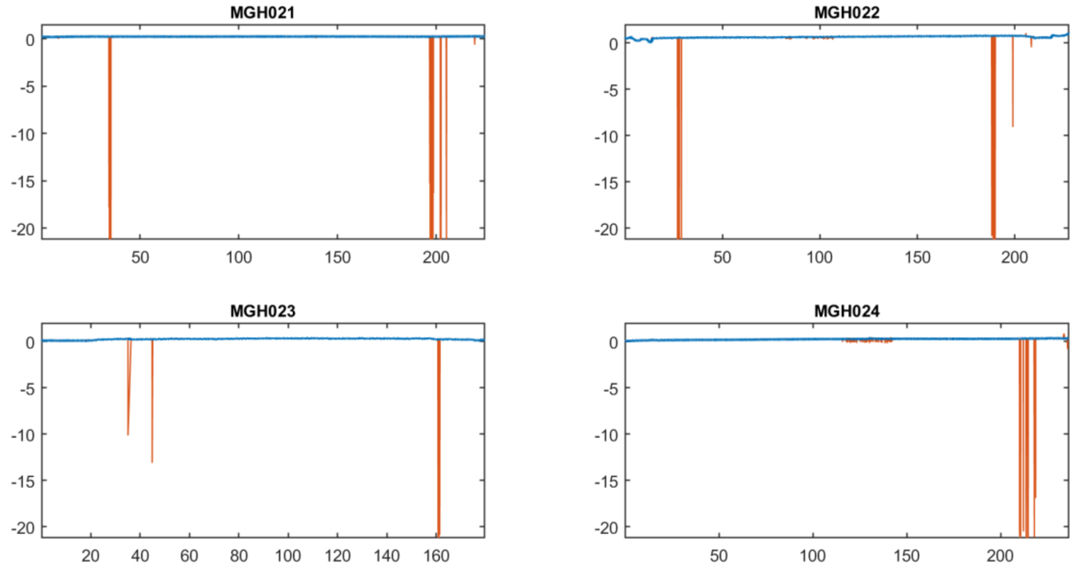


**Figure S6:** Raw and artifact-free EDA data for Subjects 21-24 collected while undergoing surgery at Massachusetts General Hospital (MGH). The raw data is in orange and the artifact-free data is in blue.

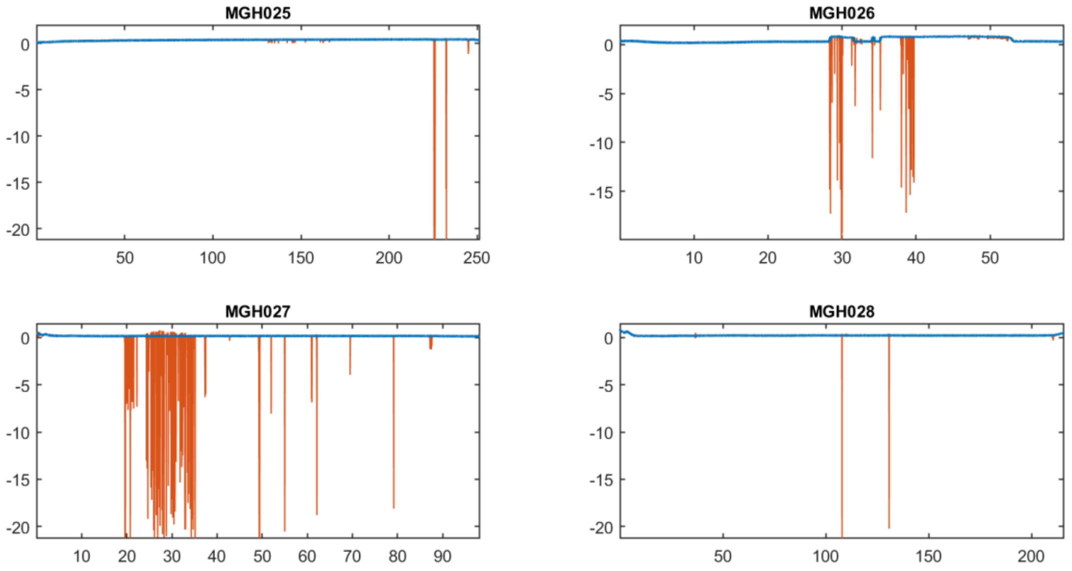


**Figure S7:** Raw and artifact-free EDA data for Subjects 25-28 collected while undergoing surgery at Massachusetts General Hospital (MGH). The raw data is in orange and the artifact-free data is in blue.

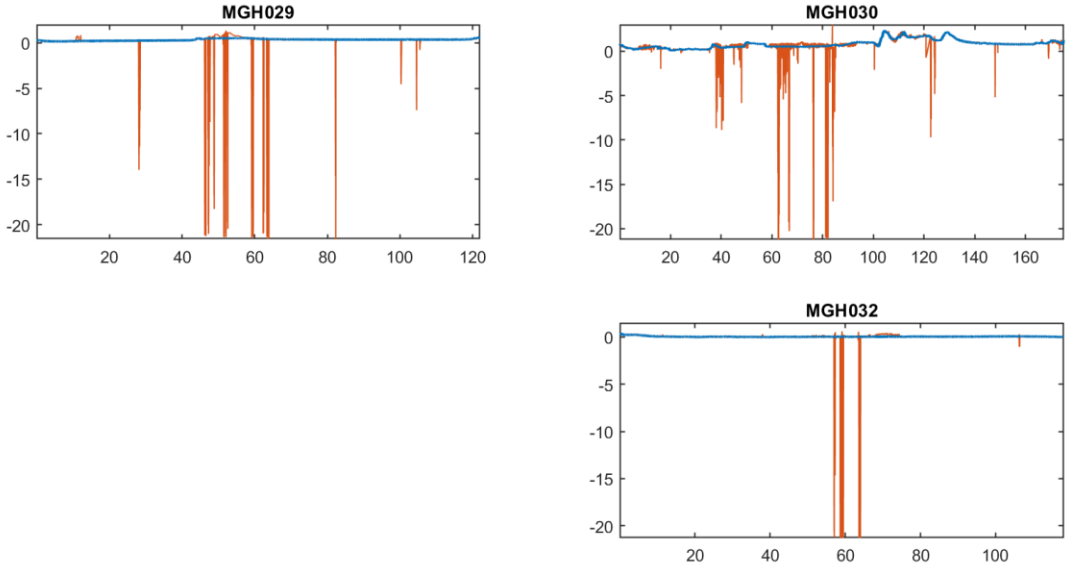


**Figure S8:** Raw and artifact-free EDA data for Subjects 29-32 collected while undergoing surgery at Massachusetts General Hospital (MGH). The raw data is in orange and the artifact-free data is in blue.

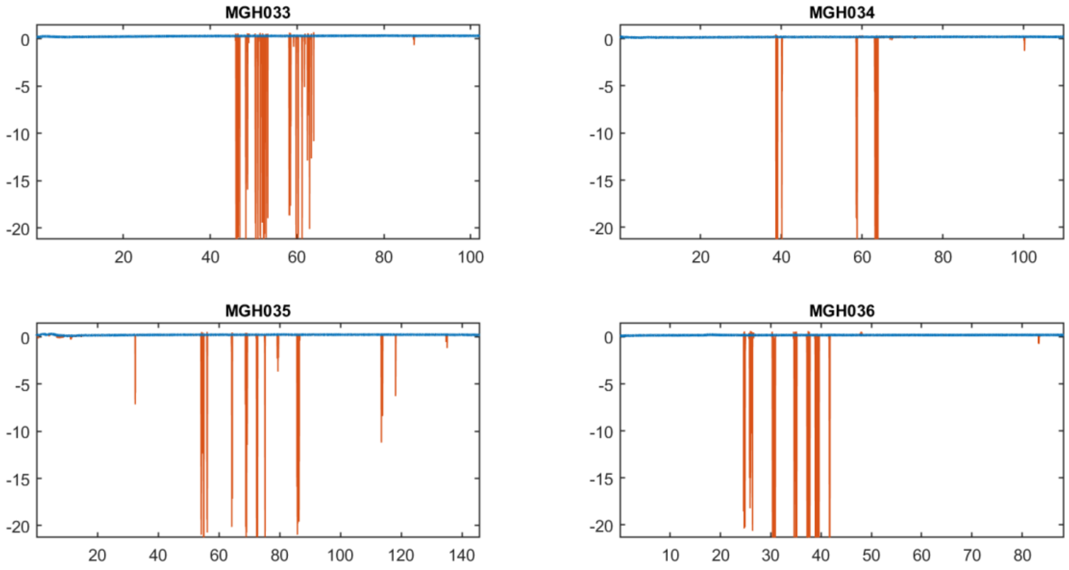


**Figure S9:** Raw and artifact-free EDA data for Subjects 33-36 collected while undergoing surgery at Massachusetts General Hospital (MGH). The raw data is in orange and the artifact-free data is in blue.

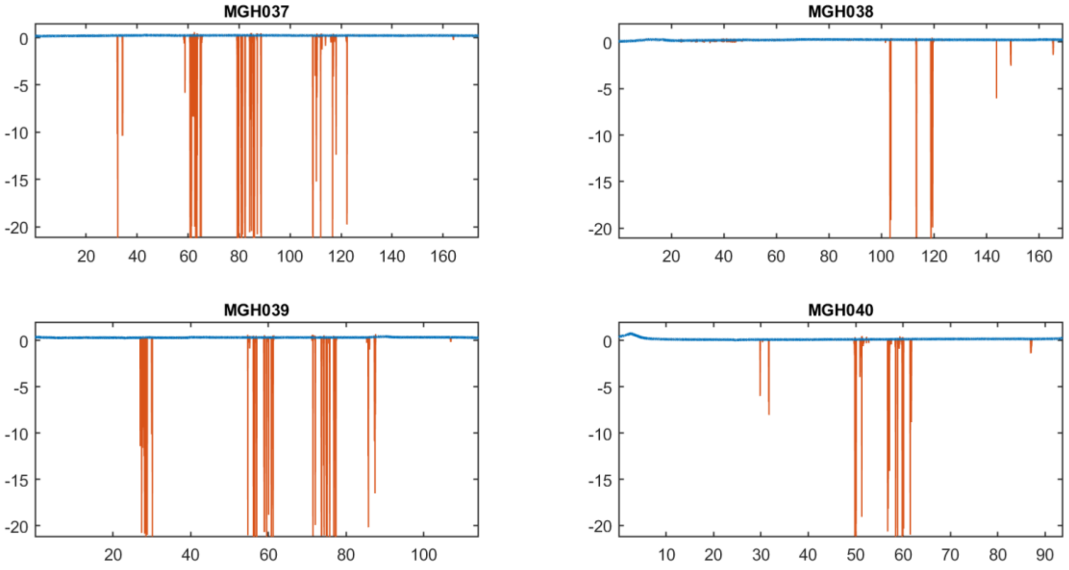


**Figure S10:** Raw and artifact-free EDA data for Subjects 37-40 collected while undergoing surgery at Massachusetts General Hospital (MGH). The raw data is in orange and the artifact-free data is in blue.

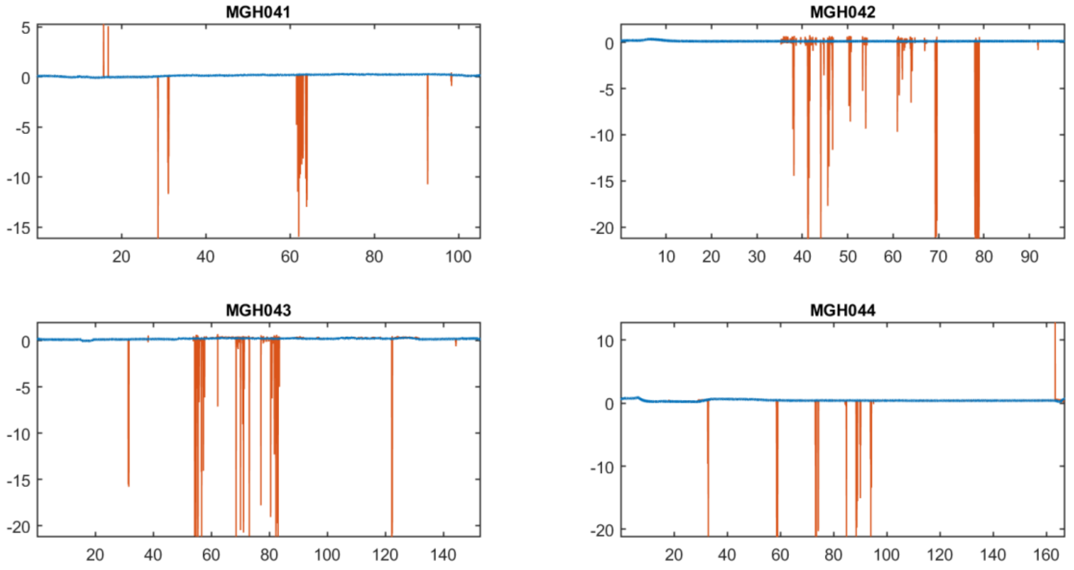


**Figure S11:** Raw and artifact-free EDA data for Subjects 41-44 collected while undergoing surgery at Massachusetts General Hospital (MGH). The raw data is in orange and the artifact-free data is in blue.

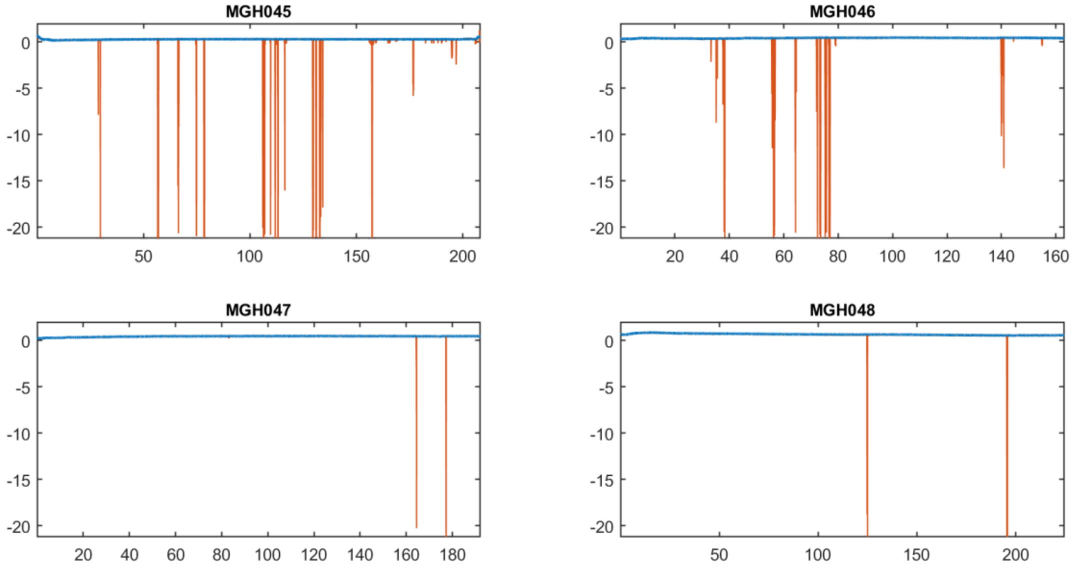


**Figure S12:** Raw and artifact-free EDA data for Subjects 45-48 collected while undergoing surgery at Massachusetts General Hospital (MGH). The raw data is in orange and the artifact-free data is in blue.

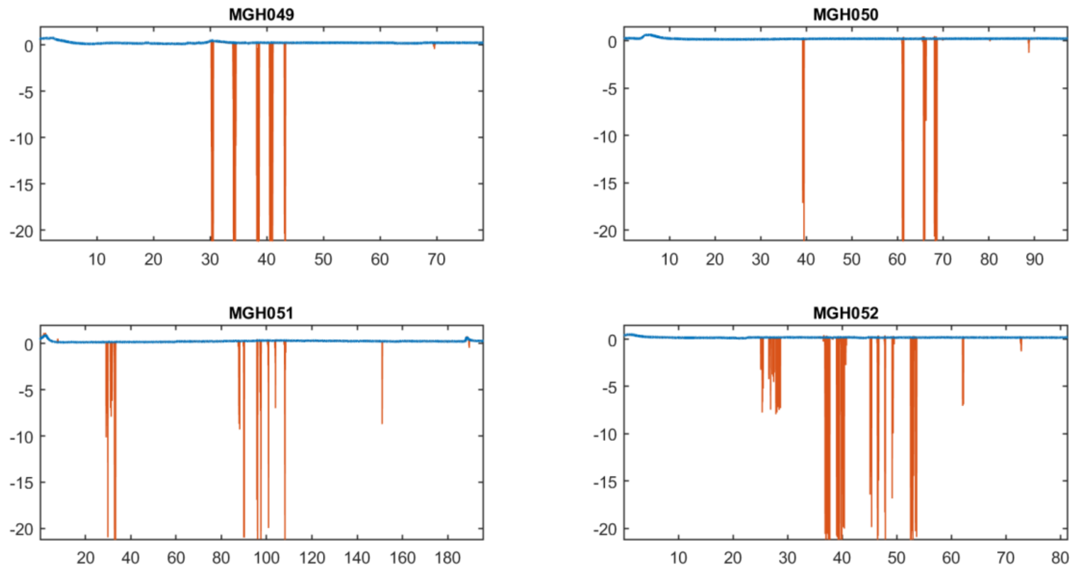


**Figure S13:** Raw and artifact-free EDA data for Subjects 49-52 collected while undergoing surgery at Massachusetts General Hospital (MGH). The raw data is in orange and the artifact-free data is in blue.

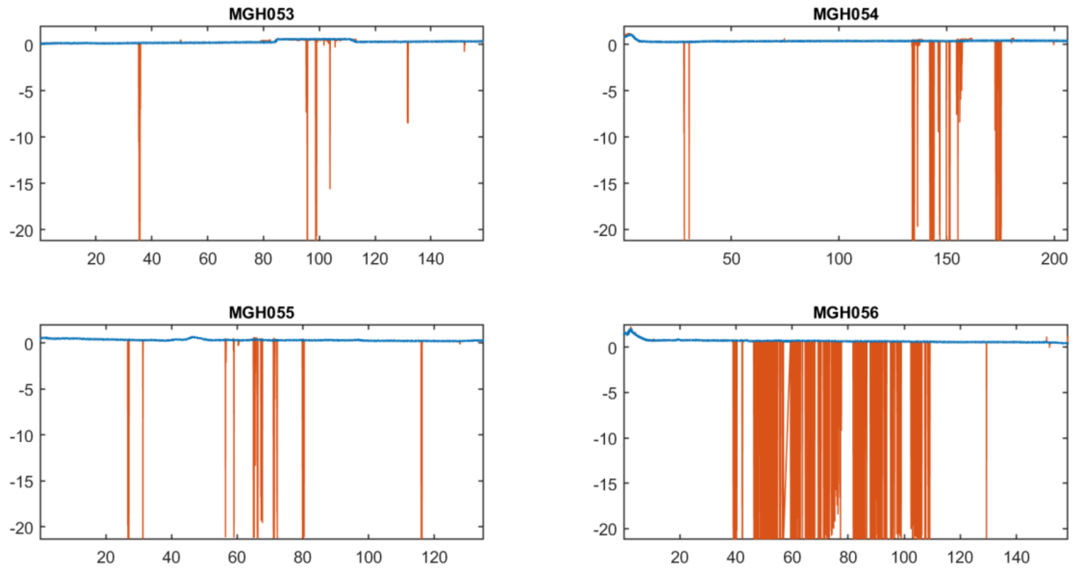


**Figure S14:** Raw and artifact-free EDA data for Subjects 53-56 collected while undergoing surgery at Massachusetts General Hospital (MGH). The raw data is in orange and the artifact-free data is in blue.

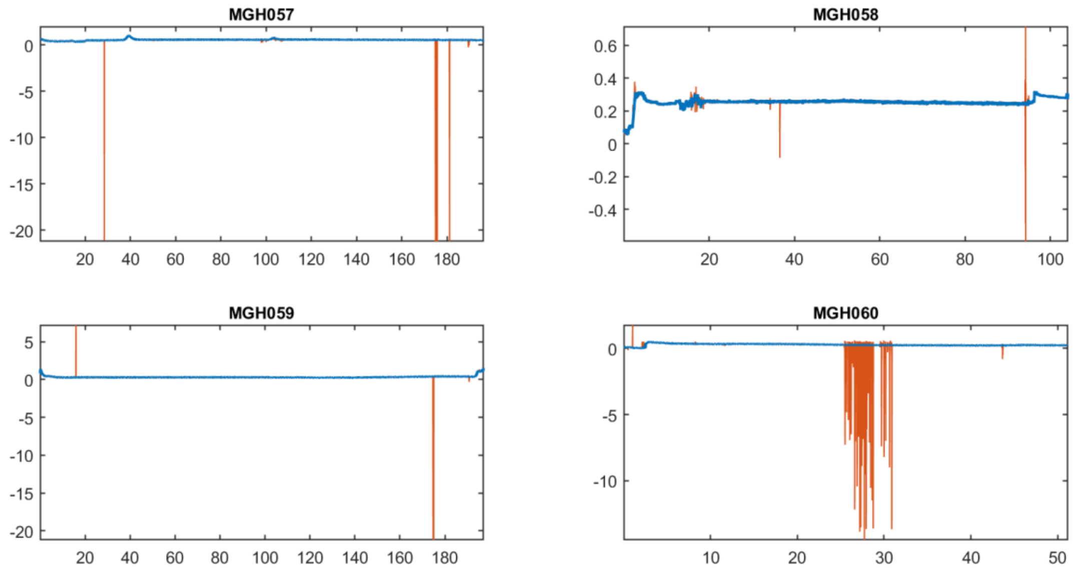


**Figure S15:** Raw and artifact-free EDA data for Subjects 57-60 collected while undergoing surgery at Massachusetts General Hospital (MGH). The raw data is in orange and the artifact-free data is in blue.

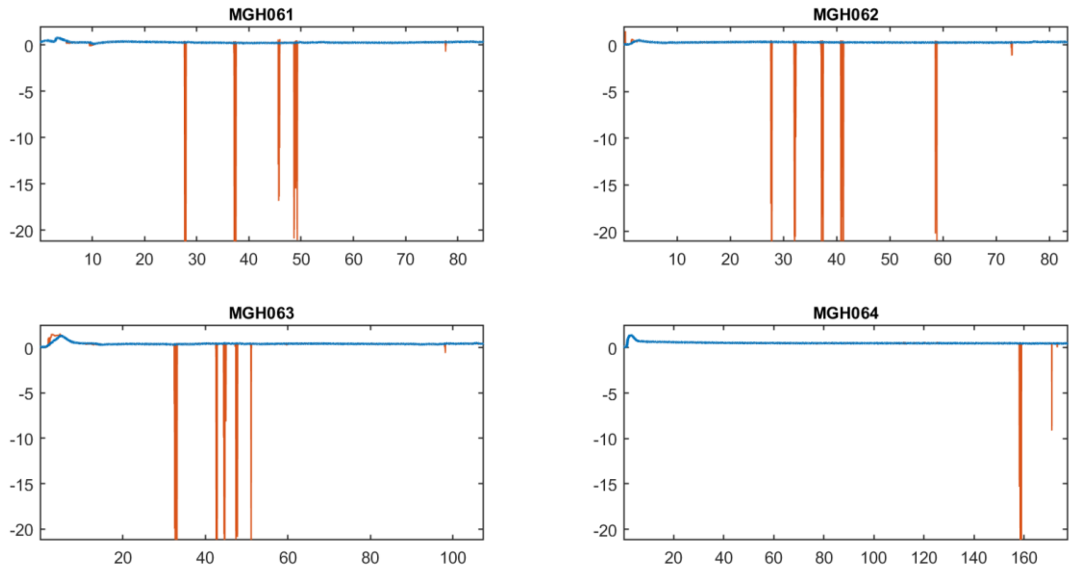


**Figure S16:** Raw and artifact-free EDA data for Subjects 61-64 collected while undergoing surgery at Massachusetts General Hospital (MGH). The raw data is in orange and the artifact-free data is in blue.

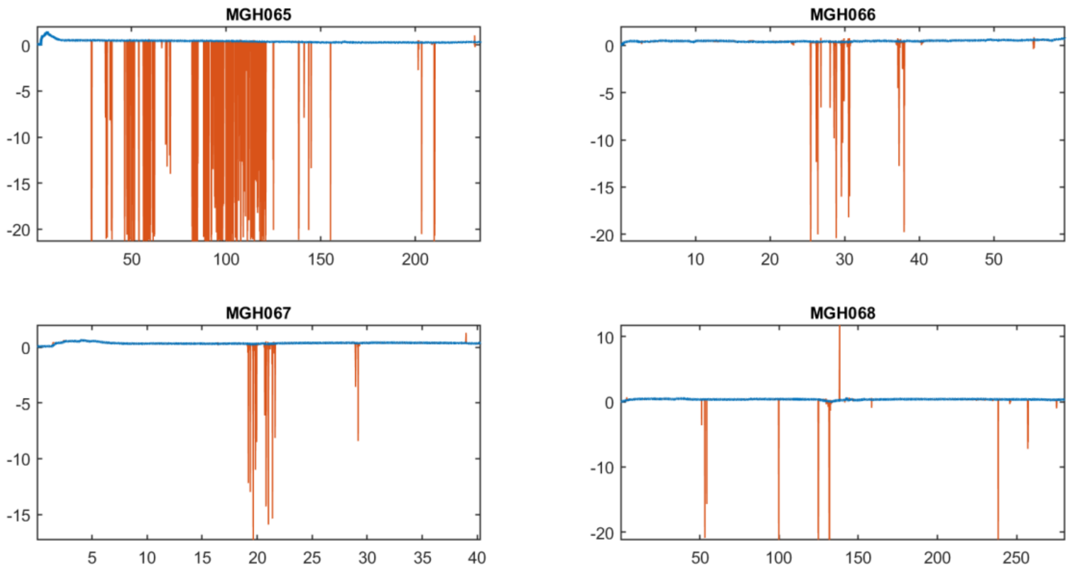


**Figure S17:** Raw and artifact-free EDA data for Subjects 65-68 collected while undergoing surgery at Massachusetts General Hospital (MGH). The raw data is in orange and the artifact-free data is in blue.

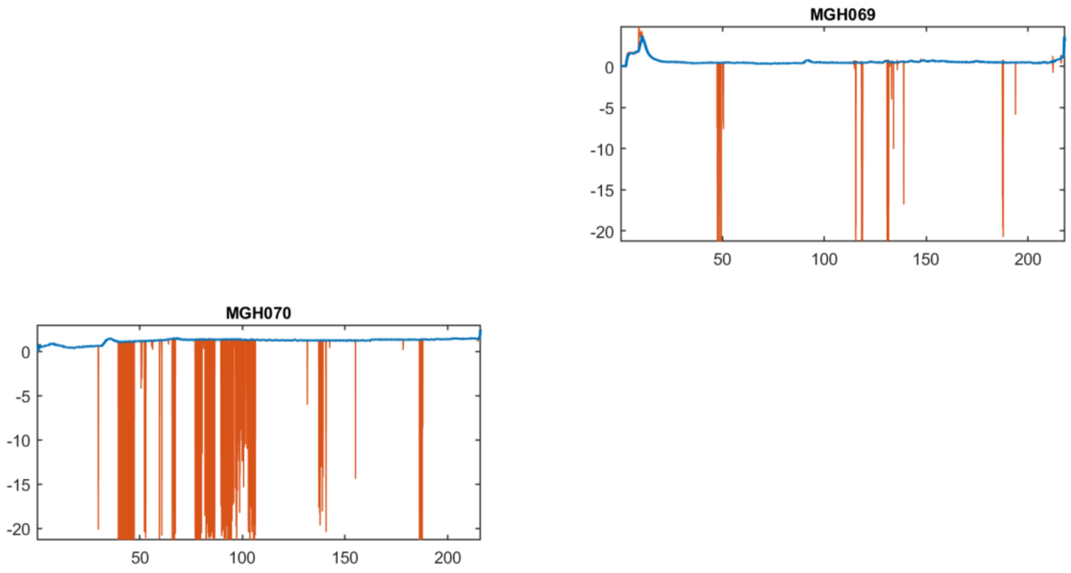


**Figure S18:** Raw and artifact-free EDA data for Subjects 69-70 collected while undergoing surgery at Massachusetts General Hospital (MGH). The raw data is in orange and the artifact-free data is in blue.
